# ERBB2 drives YAP activation and EMT-like processes during cardiac regeneration

**DOI:** 10.1101/2020.01.07.897199

**Authors:** Alla Aharonov, Avraham Shakked, Kfir Baruch Umansky, Alon Savidor, David Kain, Daria Lendengolts, Or-Yam Revach, Yuka Morikawa, Jixin Dong, Yishai Levin, Benjamin Geiger, James F. Martin, Eldad Tzahor

## Abstract

Cardiomyocyte (CM) loss after injury results in adverse remodelling and fibrosis, which inevitably lead to heart failure. ERBB2-Neuregulin and Hippo-YAP signaling pathways are key mediators of CM proliferation and regeneration, yet the crosstalk between these pathways is unclear. Here, we demonstrate in adult mice that transient over-expression (OE) of activated ERBB2 in CMs promotes cardiac regeneration in a heart failure model. OE CMs present an EMT-like regenerative response manifested by cytoskeletal remodelling, junction dissolution, migration, and ECM turnover. Molecularly, we identified YAP as a critical mediator of ERBB2 signaling. In OE CMs, YAP interacts with nuclear envelope and cytoskeletal components, reflecting the altered mechanic state elicited by ERBB2. Hippo-independent activating phosphorylation on YAP at S352 and S274 were enriched in OE CMs, peaking during metaphase, and viral overexpression of YAP phospho-mutants dampened the proliferative competence of OE CMs. Taken together, we demonstrate a potent ERBB2-mediated YAP mechanosensory signaling, involving EMT-like characteristics, resulting in heart regeneration.

**Highlights:**

1. ERBB2-driven regeneration of scarred hearts recapitulates core-EMT processes
2. YAP is activated and required downstream to ERBB2 signaling in CMs
3. YAP activity is mechanically driven by cytoskeleton and nuclear envelope remodeling
4. YAP S274 and S352 phosphorylation is essential for CM mitosis

## Introduction

Adult mammalian cardiomyocytes (CMs) have poor proliferative potential, resulting in virtually non-existent de-novo CM renewal after injury. The inability to replace the lost contractile units after acute myocardial infarction (MI) is paralleled by scar formation and fibrosis in the injured area (Tzahor and Poss, 2017). The aftermath of an acute MI inevitably leads to permanent loss of contractile force, which gradually progresses into heart failure (HF) and death. The diminished regenerative potential of the heart begins shortly after birth as CMs gradually stop proliferating, and switch from hyperplastic to hypertrophic growth (Li et al., 1996)(Bergmann et al., 2015).

A robust regenerative response, however, was demonstrated in adult hearts of lower vertebrates such as zebrafish (Poss et al., 2002) and even in neonatal mammalian hearts during a short time window after birth (Porrello et al., 2011)(Zhu et al., 2018). The source of newly formed CMs in the injury area was traced to pre-existing CMs, for both mammalian and lower vertebrates, and was shown to involve CM dedifferentiation and cell cycle re-entry (Porrello et al., 2011)(Jopling et al., 2010). This fact fuels the field of cardiac regeneration to investigate CM cell-cycle regulation (Leone and Engel, 2019)(Foglia and Poss, 2016)(Sadek and Olson, 2020). Although CM proliferation is a pre-requisite of cardiac regeneration, it has to be coordinated with other overarching processes. CM migration and ECM replacement during zebrafish heart regeneration outline the coordinated effort of multiple biological processes needed for successful regeneration (Itou et al., 2012)(Tahara et al., 2016).

The signaling network consisting of Neuregulin-1 (Nrg1), and its tyrosine kinase receptors ERBB4 and ERBB2 has been shown as a critical pathway for CM proliferation and function prenatally and postnatally (Gassmann et al., 1995)(Lee et al., 1995)(Meyer and Birchmeier, 1995)(Pasumarthi and Field, 2002)(Harvey et al., 2016). In mice, we have shown that ERBB2 levels in CMs decline after birth, resulting in CM cell cycle arrest and loss of cardiac regenerative potential. ERBB2 is necessary for the proliferative response of CMs induced by Neuregulin 1 (NRG1) during embryonic and neonatal stages, and sufficient to trigger a robust regenerative response following MI in adult mice (using a CM-restricted inducible overexpression of constitutively active *Erbb2* isoform, ca*Erbb2*-cOE, OE in short) (D’Uva et al., 2015). Similarly, inhibition of Erbb2 in zebrafish, disrupts CM proliferation and heart regeneration, while NRG1 stimulates hyperplasia in uninjured adult zebrafish hearts (Gemberling et al., 2015). In line with this, NRG1 promotes CM cycling only in an early restricted time frame when applied to human neonatal and juvenile CMs *in vitro* (Polizzotti et al., 2015).

The Hippo pathway is an important negative regulator of the cell cycle in CMs (Heallen et al., 2011)(Wang et al., 2018). The Hippo-YAP pathway has been investigated in cardiac biology via loss-of-function (cKO) and gain-of function approaches such as knockdown of the LATS kinase adaptor protein SAV1 (SAV1-cKO) and phospho-mutant YAP that evades negative regulation by the Hippo pathway (YAP 5SA/YAPS112A). Multiple lines of evidence point to the essential roles of YAP in CM proliferation and cardiac regeneration (Heallen et al., 2013)(Morikawa et al., 2015)(Heallen et al., 2017)(Monroe et al., 2019)(von Gise et al., 2012)(Xin et al., 2012). The effects of Hippo pathway inhibition and ERBB2 activation both result in CM dedifferentiation and proliferation as well as cardiac regeneration. Whether these two pathways act independently in cardiac regeneration or are interconnected, and precisely how, is unclear.

Mechanotransduction is the relay of mechanical stimuli to biochemical cues, transduced by the cytoskeleton. Mechanosensing processes were shown to robustly regulate YAP activity in an integrative manner with the Hippo pathway (Panciera et al., 2017)(Dupont et al., 2011) (Elosegui-Artola et al., 2017). Altered cellular shape, connectivity and adhesion, actin dynamics, and the continuum between the cytoskeleton and the nucleus via the ‘Linker of Nucleoskeleton and Cytoskeleton’ complex (LINC complex), are all implicated in mechanotransduction signaling. Numerous studies demonstrated a Hippo-independent YAP regulation, highlighting a multifaceted control of YAP, especially in the heart (Ragni et al., 2017)(Hirai et al., 2017)(Li et al., 2015).

In this study we have asked what underlies the potent regenerative outcome of transient ERBB2 signaling in adult mouse CMs. We revealed that ERBB2 impacts the CM cytoskeleton to promote EMT-like processes. This regenerative-EMT enables cardiac regeneration even in the context of a pre-existing scar, and involves ECM turnover and CM migration. YAP is activated and required downstream to ERBB2 signaling in CMs, in association with the altered cytoskeletons and nuclear envelope, reflective of mechanotransduction signaling. Additionally, YAP phosphorylation at S274 and S352 peaks at metaphase and is required for mitosis. In summary, we have characterized a novel ERBB2-YAP regenerative signaling axis in CMs.

## Results

### Delayed ERBB2 induction in an HF model triggers functional recovery, scar replacement and involves EMT-like processes in CMs

We have previously shown that transient ERBB2 activation promotes robust cardiac regeneration in acute MI model in juvenile and adult mice (D’Uva et al., 2015). In order to translate this regenerative signal into a more clinically-relevant model of heart disease, namely heart failure (HF), we began by exploring whether cardiac regeneration could be induced in the context of pre-existing injury. Using the transient ca*Erbb2* CM-specific induction system (D’Uva et al., 2015), we designed an experiment in which adult mice underwent MI and >3 weeks after, ca*Erbb2* was induced. Heart injuries were similar in OE and WT littermates at the 3-weeks time point (Figure 1A-E, Figure S1A). ca*Erbb2* was induced for 3 weeks, representing ∼ 2.5 weeks of ERBB2 expression, and was consistently characterized by ventricular wall hypertrophy and lumen shrinkage, which resulted in a transient increase in ejection fraction (Figure 1B-D green period, E, Figure S1B). During ca*Erbb2* induction, CMs undergo dedifferentiation, proliferation and hypertrophy (D’Uva et al., 2015) resulting in lower stroke volume and cardiac output values compared to WT mice (Figure S1C,D). Cessation of ERBB2 signaling by Doxycyclin (Dox) re-administration resulted in hypertrophy reversal and CM redifferentiation that lead to sustained improvement in numerous cardiac functional parameters (Figure 1B-E, Figure S1B-D). In addition, functional improvement was accompanied by reduced scarring in OE hearts compared to WTs (Figure 1F,G). Hence, delayed transient ERBB2 activation in CMs can trigger functional and structural regeneration in a mouse HF model.

**Figure 1:**
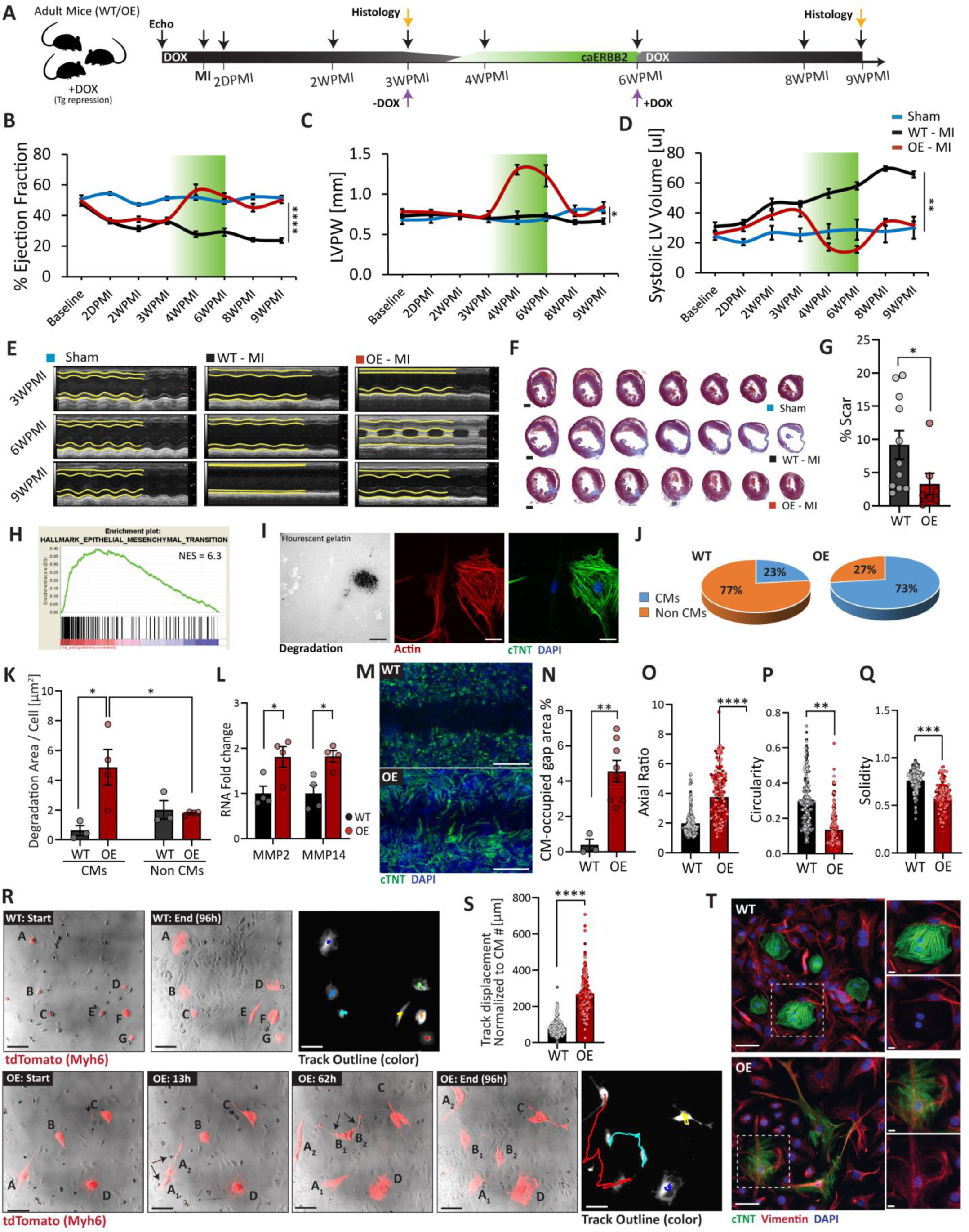
Delayed ERBB2 activation in CMs triggers functional recovery, scar replacement and involves EMT-like processes. (A) A diagram describing the experimental outline in adult mice of transient ca*Erbb2* over expression. Mice were subjected to LAD ligation (Myocardial infarction, MI), and caErbb2 induction was started 3weeks post MI (3WPMI), up to 6WPMI. Purple arrows indicate removal and re-instatement of DOX diet resulting in approximately 2.5weeks period of caErbb2 expression denoted as a green bar. Black arrows indicate Echo measurements. Orange arrows indicate histological analysis. (B-D) graphic representation of cardiac parameters derived from echocardiographic imaging analysis, performed at the indicated time points in (A). n=16 mice for WT, n=11 mice for OE, n=5 for Sham; (B) Cardiac ejection fraction %. (C) left ventricular posterior wall (LVPW) thickness. (D) Systolic left ventricular (LV) volume. (E) Representative M-mode images of anterior and posterior wall contractility for Sham, WT-MI and OE-MI, at the indicated time points. (F) Representative transverse histological sections of Sham, WT and OE hearts 9 weeks after injury stained for Masson’s trichrome showing scarring in blue. Scale bars, 1 mm. (G) Scar quantification (%) for data in (F). n=11 mice for WT, n=7 for OE. Significance was determined by one-tailed t-test. (H) Bulk RNA - seq WT-MI and OE-MI analysed using GSEA hallmark module. Representative image of gene contribution to epithelial to mesenchymal transition hallmark. (I) Immunofluorescence (IF) of indicated proteins showing degradation (black, loss of signal) in the gelatin channel co-localizing to a CM (positive for cTNT) in the perinuclear area. Scale bars, 25 µm. (J) Analysis of the proportion of degradation area in P7 cardiac cultures divided between CMs and non-CMs for OE and WT P7 cardiac cultures. (K) The specific degradation area normalized per cell for CMs and non-CMs in OE and WT P7 cardiac cultures. (n=56 CM, n=676 non-CM for WT, n=91 CMs n=680 non-CMs for OE). (L) qRT-PCR analysis of *MMP2* and *MMP14* transcripts in OE-MI and WT-MI adult heart lysates. (n=4 for WT, and n=4 for OE). (M) IF staining for the indicted proteins in a migration assay for P7 WT and OE cardiac cultures 4 days after barrier removal. Scale bars, 500 µm. (N) Quantification of CM-occupied gap area % in (M) (n=3 for WT, n=8 for OE). (O) Axial ratio quantification of OE and WT P7 CMs as a proxy for morphological changes (elongation in a particular axis) (n= 183 for WT, n=131 for OE). (P) Circularity quantification of OE and WT P7 CMs as a proxy for morphological changes. (n= 183 for WT, n=131 for OE). (Q) Solidity quantification of OE and WT P7 CMs as a proxy for morphological changes (protrusions and indentations). (n= 183 for WT, n=131 for OE). Bars are comprised of colour coded dots that differentiate data points derived from different biological repeats for (O-Q). (R) Representative images for the migration movies for WT and OE P7 cardiac cultures with CMs tagged with endogenous tdTomato fluorescent protein. The timing of each picture is denoted, arrows indicate mitosis and letters indicate specific cells throughout the images. Track outline indicates in color the individual displacement of a cell during the movie. Scale bars, 50 µm. (S) Track displacement normalized to CM number that make up a particular track in (R) plotted for WT and OE CMs. Bars are comprised of color coded dots that differentiate data points derived from different biological repeats. (n =190 for WT and n=175 for OE). (T) IF staining for the indicated proteins in P7 cardiac cultures of WT/OE. Scale bars, 50 µm. Insets are enlarged to the right. Inset Scale bars, 10 µm. \**p* < 0.05; \*\**p* < 0.01; \*\*\**p* < 0.001; \*\*\*\**p*<0.0001. Error bars indicate SEM. All experiments were performed for at least 3 biological repeats.

To begin investigating the underlying cellular characteristics of OE CMs, we performed bulk RNA - seq comparing injured WT and OE hearts. We analyzed enriched hallmark gene sets using GSEA, which highlighted epithelial to mesenchymal transition (EMT) with the highest enrichment score (Figure 1H, Figure S2A) among other hallmarks previously implicated in cardiac regeneration, such as hypoxia (Kimura et al., 2015)(Nakada et al., 2016), angiogenesis (Marín-juez et al., 2016)(Das et al., 2019) and glycolysis (Honkoop et al., 2019)(Fukuda et al., 2019). Although the myocardium is not an epithelial tissue, EMT-like behaviors such as ECM-deposition and degradation, and CM migration are in line with the requirements for tissue replacement and regeneration in the context of a pre-existing scar. We therefore set-out to probe several EMT-like processes in OE vs WT CMs. First, we tested ECM degradation using a functional fluorescent gelatin degradation assay in P7 cardiac cultures (Revach et al., 2015). Quantification revealed a strong degradation activity of the OE CMs compared to WTs (Figure 1I,J,K). RNA expression of the major EMT-related MMPs, *MMP2* and *MMP14* was elevated in OE adult hearts compared to WTs (Figure 1L).

Examination of EMT cluster gene transcripts of adult hearts revealed a profound change in overall ECM composition (Figure S2B). For example, ECM molecules reported to be enriched in P1 neonatal hearts compared to P7 (Bassat et al., 2017), such as *Col12a1*, *Col4a2*, *Col6a2*, *Col3a1* were more prevalent in OE hearts, suggestive of ECM rejuvenation. Likewise, ECM components associated with migration such as *Tnc*, *Mgp*, *Vcan*, *Postn* (Nagaharu et al., 2011)(Mertsch et al., 2009)(Inai et al., 2013)(Li et al., 2010) were enriched in OE hearts (Figure S2B). Other ECM modulators such as *Loxl2*, *Fap*, *Pcolce*, *MMP14* and *MMP2*, were also upregulated in OE compared to WT hearts (Figure S2C). *Snail* and *Twist*, the classical EMT transcription factors (Figure S2D), receptors and transmembranal proteins associated with migration and ECM-interactions such as *Itga5*, *Itgav*, *Itgb5*, *Cd44*, as well as secreted molecules involved in mammalian and zebrafish heart regeneration, *Fstl1* and *Tgfb1* respectively, were all elevated in OE hearts (Figure S2E).

In line with EMT characteristics, IF staining of cardiac tissue sections revealed loss of CM connectivity and directionality in OE hearts, compared to the organized and tightly defined WT architecture (Figure S2F). We next investigated CM motility in P7 cardiac culture migration assay. OE CMs migrated into the gap area, possessing elongated morphology and remarkable cytoskeletal protrusions, unlike WT CMs that failed to move (Figure 1M-Q). To deepen this investigation, we utilized time-lapse microscopy imaging on fluorescently-tagged OE/WT CMs with tdTomato (Movie S1,Movie S2). Single-cell CM tracking revealed that OE CMs were more motile (and proliferative) (Figure 1R,S). Staining for the mesenchymal intermediate filament protein vimentin, which is associated with migration (Liu et al., 2015), revealed that OE CMs uniquely developed a vimentin-rich cytoskeleton which is distinct from the contractile acto-myosin sarcomere typical for WT CMs (Figure 1T). Taken together, these findings demonstrate functional cardiac improvement following myocardial ERBB2 activation, in a pre-existing scar setting (aka HF model). CMs of regenerative OE hearts were characterized by EMT-like features such as migration and pronounced ECM remodeling during ERBB2 activation.

### YAP is activated downstream to ERBB2 signaling in CMs

To get a deeper insight into the mechanism of ERBB2-mediated regeneration we employed proteome and phospho-proteome analyses followed by mass-spectrometry, in addition to the RNA - seq (Figure 2A). Evaluation of predicted molecular regulators based on RNA - seq results highlighted ERBB2 and ERK which corroborate the model and previous data (D’Uva et al., 2015) (Figure 2B). The involvement of cell cycle regulators (activation of Cyclin D1 and E2f, concomitant with the inhibition of Rb1, Pten, Cdkn1a (aka P21), and Cdkn2a), is consistent with the proliferative nature of OE CMs. Interestingly, *Yap* and *Ctgf* (a target of YAP) were highlighted as upstream regulators in the OE hearts (Figure 2B). Clustering of differentially phosphorylated proteins into signaling networks identified several pathways, including actin cytoskeleton, ERK, RHOA, and Hippo. (Figure 2C). Interestingly, actin dynamics and RHOA have been linked to YAP activation via mechanosensory mechanisms (Dupont et al., 2011)(Aragona et al., 2013) while Hippo signaling is a potent canonical negative regulator of YAP (Zhao et al., 2007)(Zhao et al., 2010). Clustering of differentially expressed proteins by Biological Process GO terms, highlighted terms associated with mitosis and heart development, further underscoring regenerative aspects, in addition to emphasis on actomyosin organization and nuclear structural and regulatory features (Figure 2D).

**Figure 2:**
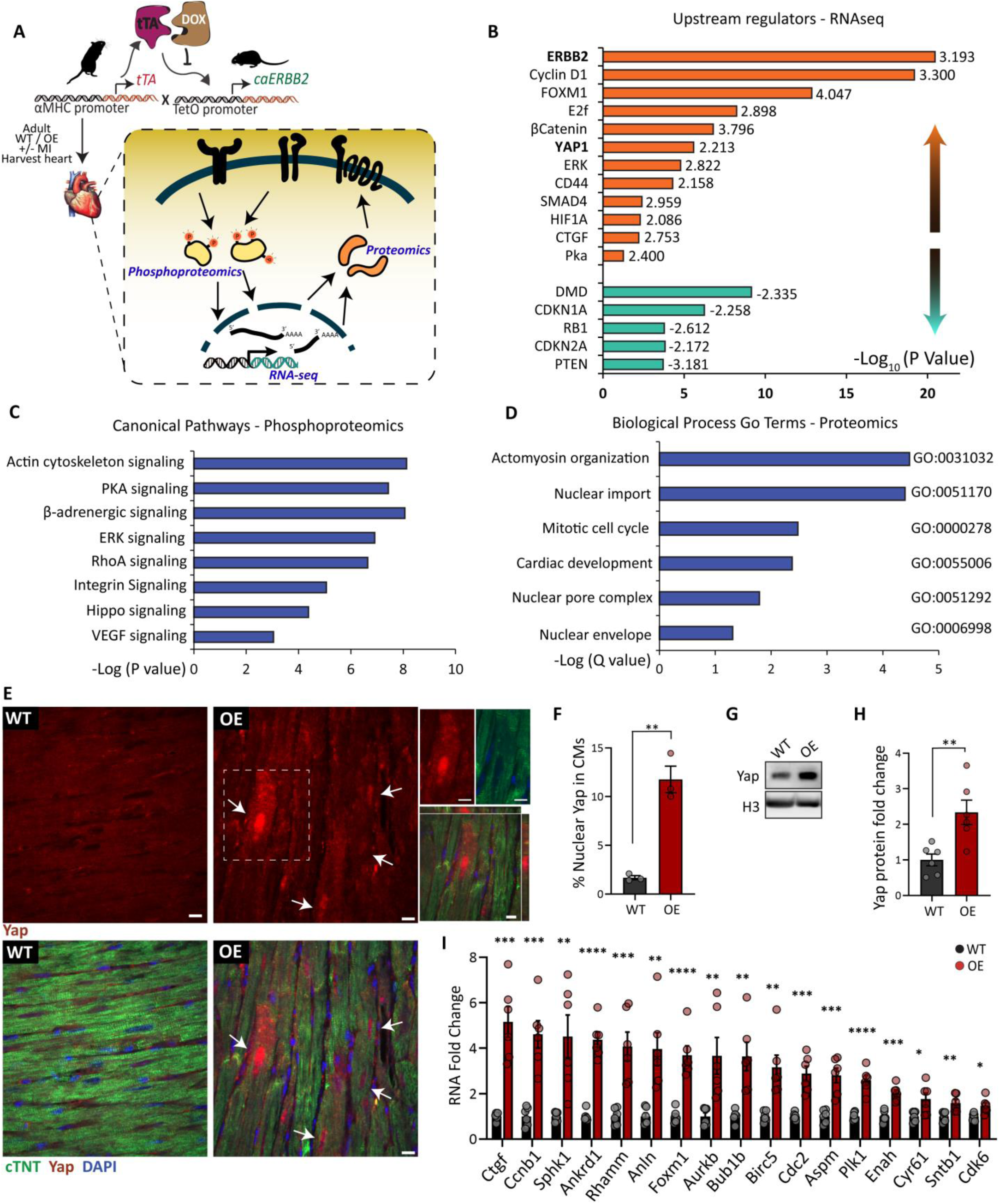
YAP is activated downstream to ERBB2 signaling in CMs. A Scheme representing the genetic model, experimental setup and assays employed. Hearts of adult WT and OE (Sham/ MI) were harvested 2 weeks after DOX removal. RNA and protein samples were simultaneously collected for RNA-seq, proteomics and phosphoproteomics analysis. n=4 for WT Sham, n=4 for WT-MI, n=4 for OE Sham, n=4 for OE-MI. (B) Predicted upstream regulators (IPA software) of differentially expressed genes between OE-MI and WT-MI using a threshold FC≥1.5, p adjusted ≤0.05. Indicated number to the right is z-score for activation (>2) or inhibition (<2). (C) Clustering of differentially phosphorylated proteins (p-value ≤ 0.05) between OE-MI and WT-MI to canonical pathways (regardless of phosphorylation) analyzed by IPA. (D) GO Clustering of differentially expressed proteins for biological processes category using PANTHER software. Marks next to bar represent GO term codes. (E) IF of indicated proteins in WT/OE heart sections. Arrows point to nuclear accumulation of Yap. Scale bar, 10 µm. Upper insets show a disrupted cTNT pattern concomitant with Yap cytoskeletal presence. Lower inset shows a z-stack image of Yap nuclear penetration. Scale bars, 10 µm. Images were taken in and around the border zone. (F) Quantification of IF in (E). Yap regarded as nuclear if the nuclear intensity was stronger than the cytoplasmic (n=1246 CMs for WT and n=713 CMs for OE). (G) WB analysis of Yap levels from in vivo adult WT/OE heart lysates. (H) Quantification of (G) (n=6 for WT and n=6 for OE). (I) qRT-PCR analysis of Yap target genes in OE/WT hearts (n=6 for WT, n=6 for OE). \**p* < 0.05; \*\**p* < 0.01; \*\*\**p* < 0.001; \*\*\*\**p*<0.0001. Error bars indicate SEM. All experiments were performed for at least 3 biological repeats.

Collectively, our high throughput (HT) analyses indicate that OE hearts are characterized by a proliferative signature involving cytoskeletal rearrangement, outlining YAP and related signaling pathways as regulators, downstream to ERBB2. Considering the central role of YAP in CM proliferation (Heallen et al., 2011)(von Gise et al., 2012)(Xin et al., 2013) and cardiac regeneration (Morikawa et al., 2017)(Heallen et al., 2017)(Wang et al., 2018), we sought to validate its involvement, and further explore ERBB2-YAP signal relay. Staining of heart sections revealed an enrichment in YAP nuclear accumulation in OE CMs (Figure 2E,F), and overall elevated YAP levels in OE CMs as shown by WB analysis (Figure 2G,H). In line with this, qRT-PCR analysis of YAP target genes revealed robust YAP transcriptional signature in OE heart lysates (Figure 2I).

### No evidence for Hippo pathway attenuation in OE hearts

To evaluate Hippo pathway activity, we analyzed the phosphoproteomic dataset obtained from WT and OE heart lysates. Quantitative phosphoproteomics indicate increased Hippo activity in OE heart lysates compared to WTs, concomitantly with proteomic data showing increased YAP protein levels, and YAP targets CTGF and ANKRD1 (Figure S3A-C). Specifically, phosphorylation sites for Hippo pathway components MOB1A and MST2 were detected on T35, and S316, respectively. T35 phosphorylation on MOB1A is a known MST site, stimulating MOB1A function (Ni et al., 2015)(Figure S3A,B). YAP S149 and S94, 2 of the 5 known LATS sites (Zhao et al., 2007) were also detected as well as the canonic LATS site S89 on Taz (Figure S3A,B). Validation of Hippo components by WB revealed higher levels of LATS1 and LATS2 kinases, MOB1 protein and phospho-MOB1 T35, as well as 14-3-3 sequestering protein in OE heart lysates (Figure S3D,E). In addition, the major negative phosphorylation site by LATS, YAP S112 (in mice, S127 in human) was also elevated in OE hearts, proportionally to YAP (Figure S3F,G). Taken together, these results do not suggest Hippo pathway silencing as a mechanism for the elevated YAP activity in ERBB2-OE hearts. Our interpretation of the data is that the active Hippo-signature acts as a negative feedback mechanism to restrain YAP activity (Moroishi et al., 2015).

### YAP is required for ERBB2-related cardiac phenotypes

We went on to evaluate the requirement of YAP to the ERBB2-OE heart phenotype by creating a transgenic mouse of both ca*Erbb2* OE and *Yap* KO with independently-activated Tamoxifen (Tam) and Dox modules that are both CM-restricted and temporally controlled (Figure 3A; mice that were both ca*Erbb2* OE and underwent Cre-mediated *Yap* excision termed OE-knockdown (OE-KD). We initiated conditional *Yap* deletion by 7 consecutive Tam injections, and after another week we induced ca*Erbb2* by Dox removal for an additional week (Figure 3A). In order to determine whether YAP is required for ERBB2 signaling, we compared the following 3 groups; WT, OE and OE-KD. The cardiac YAP KD was evaluated by expression of YAP target genes and YAP protein (Figure 3B,C), showing reduced YAP protein and transcriptional signature levels. Echocardiographic analysis revealed that ERBB2-driven elevation of ejection fraction, myocardial wall thickening, and reduced systolic and diastolic volumes, were all blunted in YAP OE-KD hearts, resulting in overall cardiac measurements between WT and OE hearts (Figure 3D-H). The proliferation markers Ki67 and PH3 were reduced in OE-KD compared to OE CMs, indicating that YAP is required for ERBB2-mediated CM proliferation. The expression of prominent EMT markers (Lamouille et al., 2014) such as *Snail* and *Twist*, *fibronectin*, *MMP2* and *MMP14*, were reduced in OE-KD hearts compared to the elevation seen in OE hearts (Figure 3K). Finally, histological analysis revealed reduced tissue disarray and intercellular gaps in OE-KD hearts reflecting an intermediate stage between WT and OE groups (Figure 3L). Taken together, these results demonstrate that YAP is indispensable for ERBB2 signaling outcomes in CMs.

**Figure 3:**
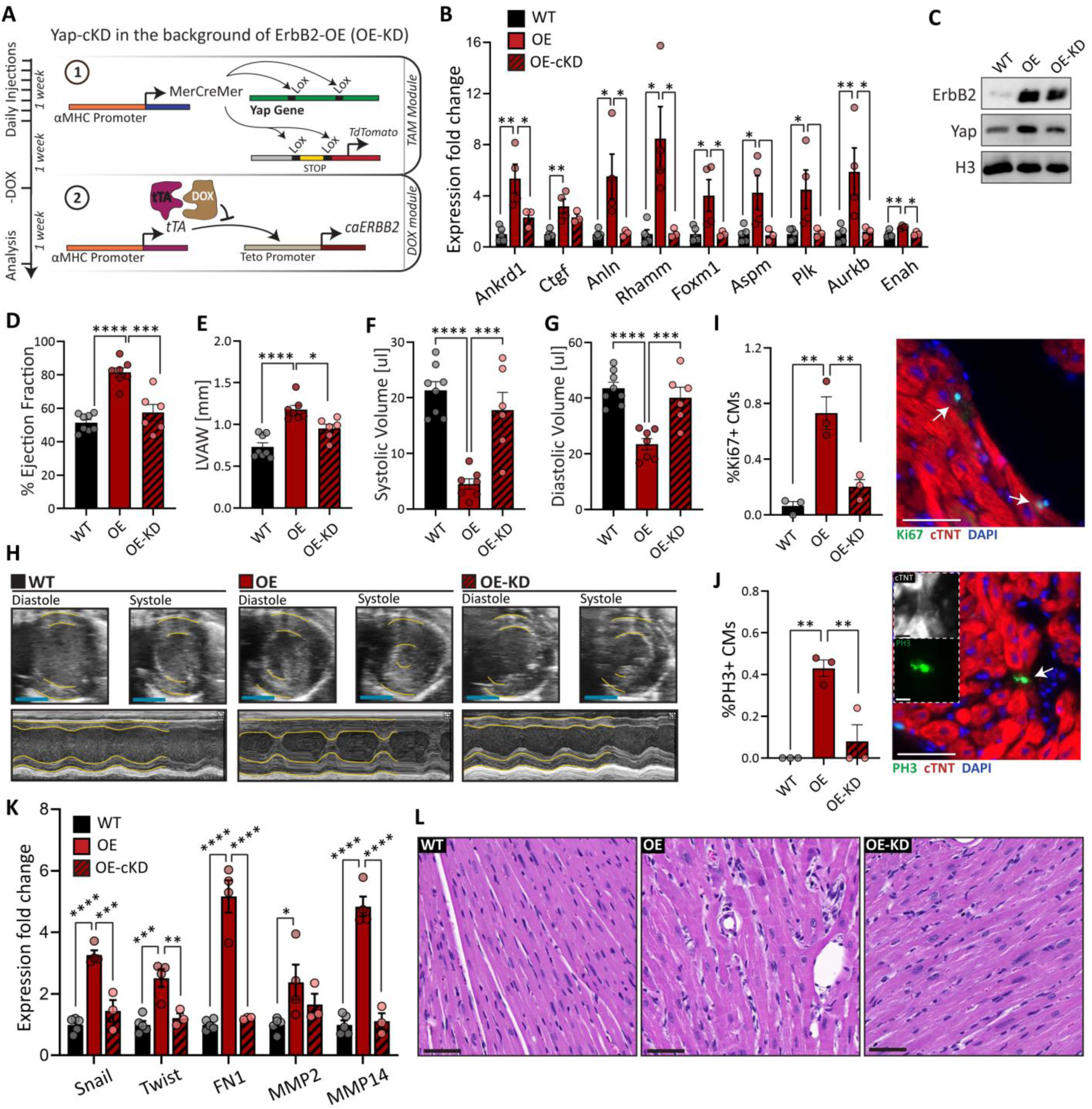
Yap is required for ERBB2-related cardiac phenotypes. (A) Schematic illustration of the OE-KD mouse model and induction regimes for Tamoxifen (TAM) and doxycycline (DOX), controlling Yap deletion and caErbB2 over expression, respectively. (B) qRT-PCR analysis of Yap target genes in WT, OE and OE-KD heart lysates (n=5 for WT, n=4 for OE, and n=3 for OE-KD). (C) WB analysis of the indicated proteins from in vivo adult WT, OE and OE-KD heart lysates. (D-G) cardiac parameters acquired from echocardiography analysis, comparing WT, OE and OE-KD mice. n=8 mice for WT, n=7 mice for OE, n=6 for OE-KD; (D) Ejection fraction %; (E) Left ventricular anterior wall (LVAW) Thickness; (F) Systolic volume (G) Diastolic volume; (H) M-mode representative images of heart in diastole and systole. Yellow tracing reflects of anterior and posterior wall contractility, for WT, OE and OE-KD mice. (I) IF analysis of % Ki67 positive CMs for WT, OE and OE-KD heart sections. Scale bar 50 µm. Arrows point to Ki67^+^ CM. (J) IF analysis of %PH3 positive CMs for WT, OE and OE-KD heart sections. Scale bar 50 µm. Arrow points to PH3^+^ CM (in metaphase). Insets highlight the disrupted sarcomere in the cTNT channel in grey (upper) and the condensed chromatin marked with PH3 with green (bottom). Scale bar 10 µm. (K) qRT-PCR analysis of EMT hallmark genes in WT, OE and OE-KD heart lysates (n=5 for WT, n=4 for OE, and n=3 for OE-KD). (L) Haematoxylin and Eosin stained histological sections of WT, OE and OE-KD hearts. Scale bar 50 µm. \**p* < 0.05; \*\**p* < 0.01; \*\*\**p* < 0.001; \*\*\*\**p*<0.0001. Error bars indicate SEM. All experiments were performed for at least 3 biological repeats.

### ERBB2 signaling induces an altered mechanical state in CMs

To gain insight into YAP signaling driven by ERBB2 activation, we employed an unbiased and quantitative Co-IP-MS by immunoprecipitating YAP in OE and WT heart lysates, followed by Mass-Spec analysis in order to identify YAP binding proteins (Figure 4A). YAP binding partners enriched in OE hearts were related to nuclear envelope, cytoskeletal, and nuclear proteins (Figure 4B-D; for full binding partner list see Figure S4). The majority of the nuclear interactors are histone proteins, consistent with the chromatin remodeling activity of YAP in the heart (Monroe et al., 2019), further corroborating YAP nuclear accumulation in OE CMs (Figure 4D). The mechanosensitive actin binding protein, Flnc and two of its associated proteins, Hspb7 and Xirp2, were enriched YAP binding partners in OE hearts (Figure 4C), the latter being implicated also in mechanotransduction in the inner ear (Leber et al., 2016)(Juo et al., 2016)(Scheffer et al., 2015). Further, YAP interaction with the intermediate filament protein Nestin was elevated in OE hearts. Nestin, a characteristic component of embryonic CMs, was also shown to correlate with YAP activation in fetal CMs (Hertig et al., 2018). Finally, the nuclear envelope proteins Sun1, Sun2 (both part of the LINC complex), Tor1aip1, Tmpo and LaminA/C were enriched YAP interactors in OE hearts. These proteins, implicated in nuclear mechanosensing, form a complex at the inner nuclear membrane (INM) with other LINC components (Hetzer, 2010)(Stroud et al., 2014). Given the substantial involvement of YAP mechanosensory fingerprints in the Co-IP assay, we focused on the differences in mechanical properties of OE and WT hearts, starting with the nucleus, the main mechanosensing organelle. Nuclear projected area was elevated in OE CMs mostly in the transverse axis, suggesting that OE nuclei experience more mechanical stress (Figure 4F-H)(Kirby and Lammerding, 2018). Consistently, Lamin A/C levels in OE hearts were elevated (Figure 4I), also known as “stress strengthening” positive feedback that withstands forces and protects nuclei experiencing formidable mechanical strain against fragility (Swift et al., 2013)(Cho et al. 2017). Lamin A/C phosphorylation on S22 was also elevated in OE hearts (Figure 4I). This mechanosensitive phosphorylation was shown to promote Lamin A solubilization into the nucleoplasm and is associated with mitotic NEBD (Torvaldson et al. 2015)(Ward and Kirschner, 1990)(Heald and McKeon, 1990)(Buxboim et al., 2014). Likewise, analysis of the proteome revealed an elevation in several cytoplasmic membrane-bound and nuclear-envelope bound force sensors and transducers in OE hearts (Figure 4J) (Elosegui-artola et al., 2016).

**Figure 4:**
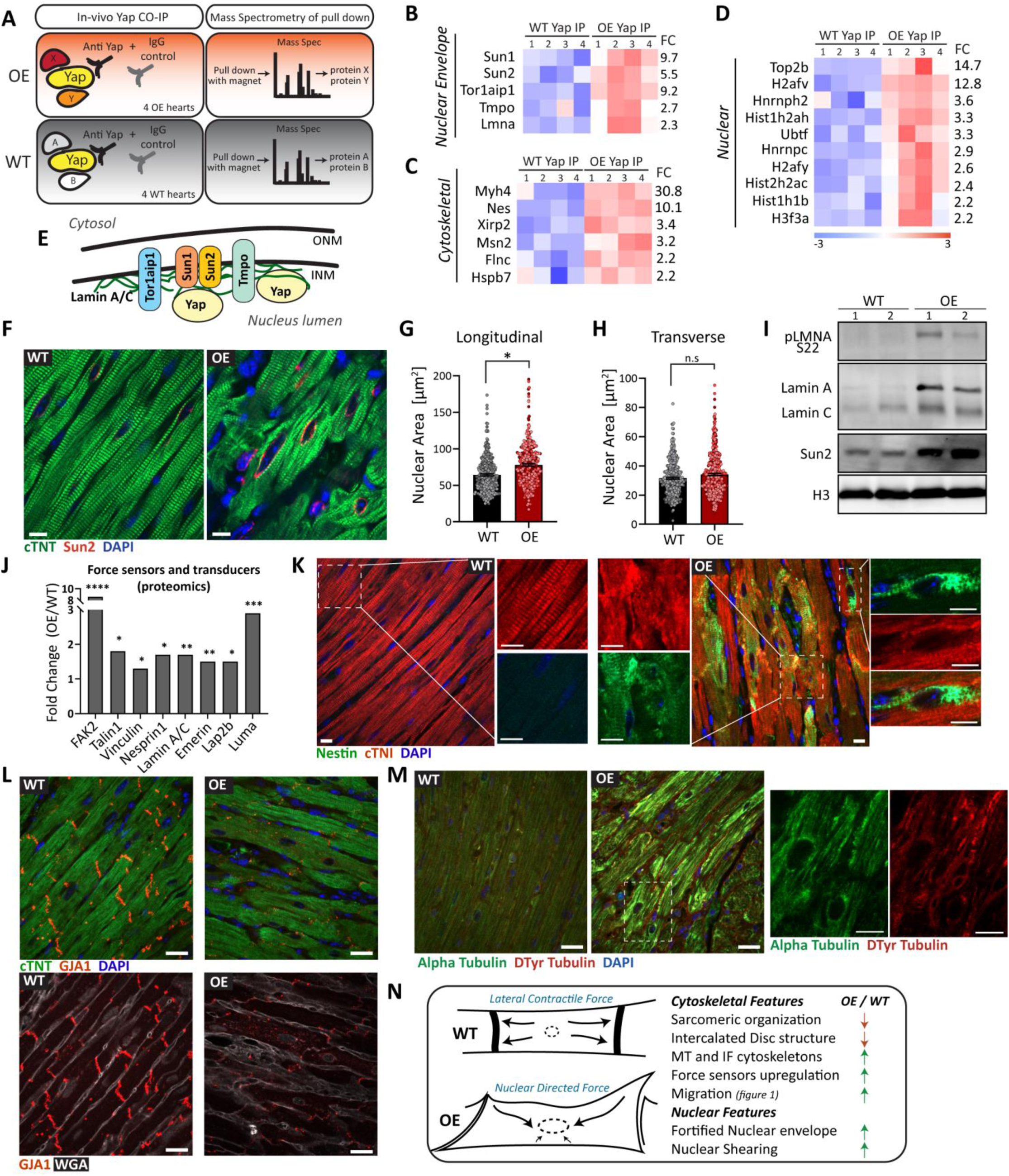
ERBB2 signaling induces an altered mechanical state in CMs. (A) Schematic illustration of Co-IP-MS experiment (n=4 IP reactions in WT, n=4 IgG reactions in WT, n=4 IP reactions in OE, n=4 IgG reactions in OE). (B) Heat map of Yap binding partners from the nuclear envelope, cytoskeleton (C), and nucleus (D). For panels (B-D) FC to the right indicates fold change of the detected protein by Mass Spec. All displayed results are at p-value ≤ 0.05. Full binding partner list appears at Figure S4. (E) Illustration of Yap associated proteins from (B) that can complex together at the inner nuclear membrane (INM). (F) IF analysis of heart sections for the indicated proteins. Scale bar 10µm. (G) Quantification of CM nuclei cross-sectional area from (F) in longitudinally cut CMs (n=313 for WT, n=361 for OE), and in transversely cut CMs (n=349 for WT, n=394 for OE) in (H). For panels (G-H) Bars are comprised of color coded dots that differentiate data points derived from different biological repeats. (I) WB analysis of the indicated protein in adult WT/OE heart lysates. (J) Proteomic data of OE/WT fold change of the indicated proteins (n=4 hearts for WT, n=4 hearts for OE). (K) IF analysis of heart sections for the indicated proteins for a WT/OE. Insets show separately two of the channels to illustrate the structural disassembly of cTNI and Nestin accumulation in OE hearts compared to WT. Scale bar 10µm. (L) IF analysis of heart sections for the indicated proteins for a WT/OE. Scale bar 10µm. (M) IF analysis of heart sections for the indicated proteins for a WT/OE. Inset of the OE image shows separately two of the channels. Scale bar 10µm. (N) Scheme summarizing differences in force distribution between a WT/OE CMs based on nuclear and cytoskeletal parameters. \**p* < 0.05; \*\**p* < 0.01; \*\*\**p* < 0.001; \*\*\*\**p*<0.0001. Error bars indicate SEM. All experiments were performed for at least 3 biological repeats.

The cytoskeleton of OE CMs was substantially enriched with Nestin at the expense of the striated sarcomeric protein cTNI. WT CMs, conversely, displayed organized, striated sarcomeres as evidenced by cTNI staining (Figure 4K, insets, Figure S5A,B). The major intercalated disk gap junction protein Gja1, which is required for proper mechanical and electrical coupling between CMs (Delmar and Liang, 2012)(Solan and Lampe, 2009) was downregulated and displaced in OE CMs (Figure 4L, Figure S5C,D). Finally, assessing the microtubule network (required for mitosis) reveled a striking increase in microtubules in OE CMs, as well as elevation of dTyr tubulin, a PTM on tubulin that hampers sarcomeric contractility (Figure 4M, Figure S5E,F) (Robison et al., 2016). Taken together, these findings indicate that all three cytoskeletal networks of OE CMs (microfilaments, intermediate filaments and microtubules) were remodeled following ERBB2 activation, thus shifting the CMs from contractility to force sensing and regulating function, and suggests the substitution of the lateral contractile force propagation in WT CMs to a nuclear directed force in OE CMs (Figure 4N).

### YAP phosphorylation on S274 and S352 peaks during metaphase, and is required for cell division

To further deepen our understanding of YAP regulation elicited by ERBB2 activation, we focused on (non-Hippo) phosphorylation sites of YAP at S274 (S289 in human) and S352 (S367 in human) detected in our phosphoproteomics and YAP IP-MS profiling. Using a custom-made antibody against pYAP S274 (Yang et al., 2013) and a newly generated custom antibody against pYAP S352, we validated the enrichment of these YAP phospho-forms in OE heart sections (Figure 5A,B). In OE adult CMs *in vivo* pYAP S274 demonstrated a diverse pattern localizing mostly to cytoskeletal compartments, but was also perinuclear and nuclear (Figure 5A), while pYAP S352 was strongly nuclear. Both pYAP S274 and pYAP S352 were present, to some extent, in intercalated discs, as shown before for general YAP (Morikawa et al., 2017)(Lin and Pu, 2015). Levels of pYAP on S274 and S352 measured by WB, similarly showed enrichment in OE heart lysates (Figure S6A,B). Interestingly, in addition to the typical YAP protein, we identified and validated a lower molecular weight isoform of YAP, which is mostly detected by antibodies to phospho-YAP S274/S352, to be enriched in OE CMs (Figure S6A-F).

**Figure 5:**
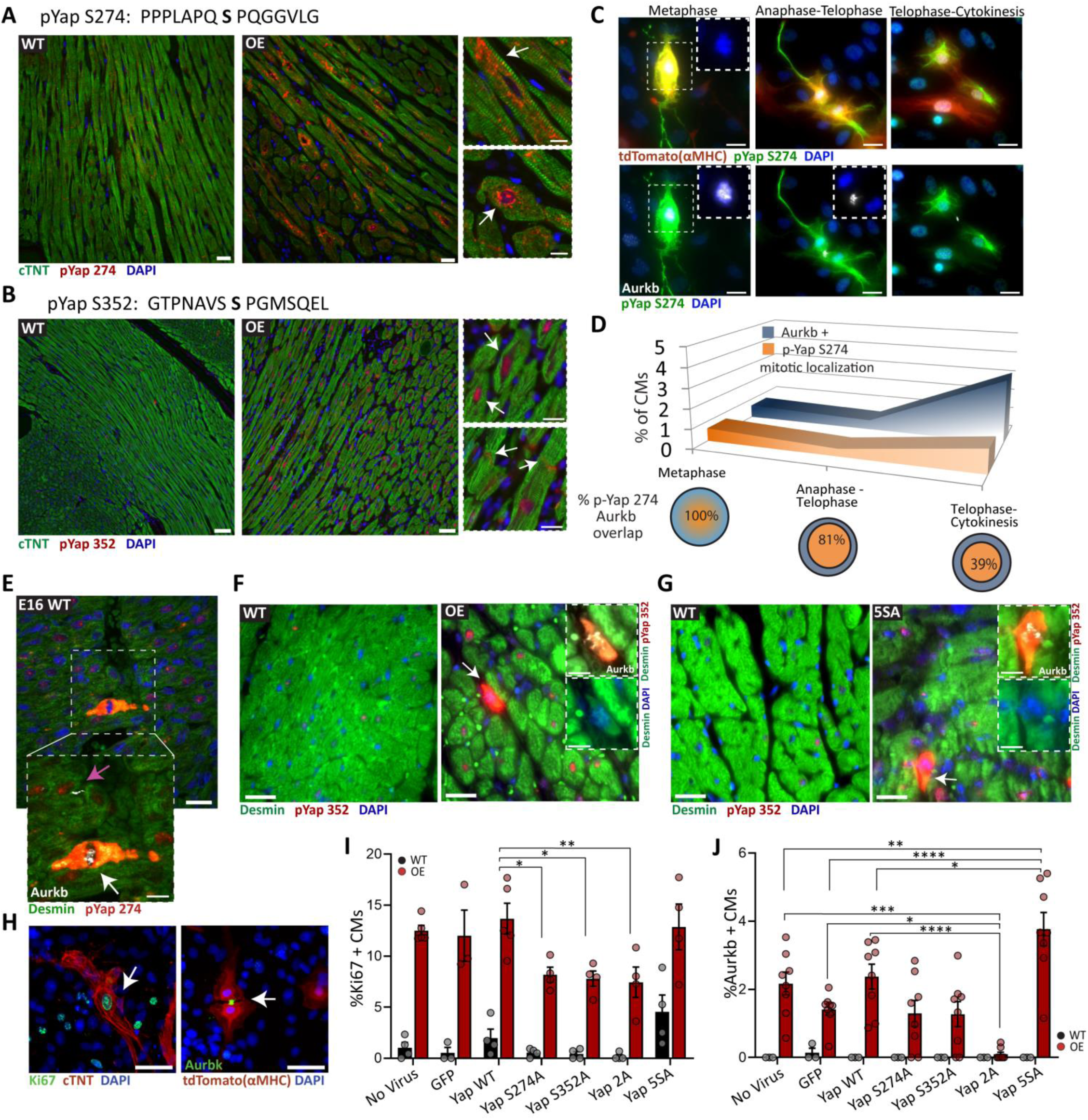
YAP phosphorylation on S274 and S352 peaks during metaphase, and is required for cell division. (A) Upper: Phosphorylation of highlighted serine on Yap identified by Yap-IP-MS (Figure 4A) enriched in OE adult heart lysates (*pv* <0.05). Lower: IF analysis of adult heart sections of the indicated proteins for WT/OE. Scale bar 20µm. Upper inset depicts the cytoskeletal morphology abundant in OE CMs, lower inset depicts the nuclear (and nuclear envelope) morphology abundant in OE CMs. Scale bar 10µm. (B) Upper: Phosphorylation of highlighted serine on Yap identified by MS phosphosproteomic analysis (Figure 2A) enriched in OE adult heart lysates (*pv* <0.05). Lower: IF analysis of adult heart sections of the indicated proteins for WT/OE. Scale bar 50µm. Upper inset depicts the nuclear accumulation abundant in OE CMs, lower inset shows the presence in intercalated discs. Scale bar 20µm. (C) Representative IF images for the indicated proteins at the metaphase, anaphase-telophase and telophase-cytokinesis stages in P7 OE cardiac cultures (CM lineage endogenously tagged with tdTomato). Scale bar 20µm. (D) Quantification of CMs positive for Aurkb (blue curve) and pYap S274 mitotic peak localization (orange curve) counted at the different stages of mitosis plotted on the x axis. Bottom: Venn diagram show the degree of overlap between of pYap S274 mitotic peak localization from all Aurkb events. (n=2327 CMs) (E) IF analysis of embryonic WT E16.5 heart sections for the indicated proteins. Lower inset shows selected area counterstained with Aurkb. White arrow points to a CM in metaphase and pink arrow points to cytokinesis. Scale bar 10µm. (F) Immunofluorescence analysis of heart sections for the indicated proteins for WT/OE. Scale bar, 25µm. OE Insets show, with different channels, a CM in metaphase. Scale bar, 10µm. (G) IF analysis of heart sections for the indicated proteins for WT/5SA. Scale bar, 25µm. 5SA Insets show, with different channels, a CM in metaphase. Scale bar, 10µm. (H-J) WT/OE P7 cardiac cultures infected with the indicated AAV viruses, and stained for cell cycle markers Ki67 and Aurkb; (H) Representative IF images of CMs positive for Ki67 and Aurbk. Scale bar, 50µm. (I) Quantification of the Ki67% positive CMs (n= 3454 CMs for WT, n=6482 CM for OE), and (J) Aurkb% positive CMs (n= 2576 CMs for WT, n=12615 CM for OE). Signal for Aurkb was considered positive from metaphase onwards. \**p* < 0.05; \*\**p* < 0.01; \*\*\**p* < 0.001; \*\*\*\**p*<0.0001. Error bars indicate SEM. All experiments were performed for at least 3 biological repeats.

IF analysis on P7 cardiac cultures with anti pYAP S274 antibody revealed that while all cultured fixed cells have some level of nuclear staining, pYAP S274 in OE CMs had a strong filamentous appearance, as well as perinuclear and nuclear, which peaked during mitosis throughout the cell (judging by chromatin condensation) (Figure S6G). Likewise, pYAP S352 staining in cardiac cultures peaks throughout the cell during mitosis, and otherwise was enriched in the nuclei of OE CMs (Figure S6H). This expression pattern recapitulates the mitosis pattern observed in non-CMs for pYAP S274 (Figure S6 G,H, insets in WT section) (Yang et al., 2013)(Bui et al., 2016). Co-staining of the mitosis marker Aurora kinase B (Aurkb) validated that pYAP S274 and S352 peak during mitosis (Figure 5C,D and Figure S6I,J). Intensified staining was seen during metaphase, during which all CMs were double positive (in existence, not localization) for Aurkb and pYAP S274 or pYAP S352. Staining for pYAP S274 and S352 further revealed 81% and 38% of Aurkb^+^ CMs during anaphase-telophase, and 39% and 18% Aurkb^+^ CMs at telophase-cytokinesis, correspondingly (Figure S6K). The latter stage took longest and therefore had proportionally more CMs at the moment of fixation (Figure 5C,D, Figure S6I,J)). In order to determine whether pYAP S274 and S352 characterize dividing CMs during normal development, we stained E16.5 embryonic WT hearts and detected a strong nuclear pYAP S274 and S352 staining that peaks during chromosome condensation. Co-staining with Aurkb demonstrated that these were CMs undergoing metaphase, and the staining was mostly diminished upon cytokinesis completion (Figure 5E, Figure S6L).

We next examined pYAP S274 expression patterns in hearts over-expressing a constitutively active form of YAP (YAP5SA) rendering it immune to inhibition by the Hippo pathway (Monroe et al., 2019). Similarly to OE hearts *in vivo*, YAP 5SA CMs displayed a peak of both pYAP S274 and S352 during metaphase (Figure 5F,G and Figure S8M), although YAP 5SA CMs lacked the prominent pYAP S274 cytoskeletal pattern observed in OE hearts. Finally, we evaluated the requirement of the two YAP phosphorylation sites, S274 and S352, for mitosis, by infecting P7 OE/WT cardiac cultures with AAV harboring serine-to-alanine YAP phosphomutants in both sites separately and together (YAP2A), and analyzed their proliferative potential. Staining for Ki67 (broad cell cycle progression) and Aurkb (mitosis), revealed a reduction in cell cycle activity for the phospho-mutants and particularly for the YAP2A mutant for ERBB2-mediated CM mitosis (Figure 5H-J). Overall, these findings demonstrate the requirement of YAP phosphorylation on S274 and S352 during CM mitosis. Apart from mitosis, the phosphorylation of YAP on S274 is strongly associated with the altered cytoskeleton in OE CMs, correlative with YAP activation.

## Discussion

This study addresses the crosstalk of two major signaling hubs in CMs, ERBB2 and YAP, and presents numerous novel insights in the field of cardiac regeneration. First, we demonstrate that even a scarred heart (i.e. HF model) is amenable to regeneration by myocardial activated ERBB2 signaling, thus broadening the therapeutic possibilities of heart failure patients, a notion that has only recently emerged in the field of cardiac regeneration (Heallen et al., 2017). Our study demonstrates, for the first time to our knowledge, core EMT-like processes during cardiac regeneration, beyond CM dedifferentiation and proliferation. The emerging massive cooperation of ECM and cytoskeletal remodeling with CM migration seems to be pivotal for scar replacement by new CMs. This is in line with other studies suggesting that CM migration is essential for cardiac regeneration (Zhang et al., 2013)(Tahara et al., 2016)(Morikawa et al., 2015).

Mature CMs are highly specialized, stationary and strongly coupled to each other, an un-intuitive cell for EMT-like behaviors. In the embryo, processes which recapitulate core-EMT behaviors are essential to proper cardiac development and morphogenesis, including progenitor migration, trabeculation and valve formation (Cortes et al., 2017)(Thiery et al., 2009)(del Monte-Nieto et al., 2018)(Liu et al., 2010)(Cherian et al., 2016)(Von Gise and Pu, 2012)(Combs and Yutzey, 2009). We show that ERBB2 activation coerces adult CMs to become more fibroblast- and embryonic-like in nature, performing competitively in the context of existing fibrosis by movement, proliferation and engagement with the ECM. The EMT-like features seen in OE CMs are accompanied by robust changes in the cytoskeletal infrastructure, which underlies their biology. This also reflects a high degree of dedifferentiation, reminiscent (at least in part) of embryonic features, (Nieto, 2013). The EMT-like biology seen following ERBB2 activation during cardiac regeneration bears striking resemblance to cancerous processes (Nieto et al., 2016), illustrating that the underlying mechanisms of malignancy and regeneration may ‘draw from the same deck’.

Concomitantly, this also highlights the importance of the transience (i.e. eventual cessation) of the oncogenic signaling for functional rebound (Gabisonia et al., 2019)(Mohamed et al., 2018). We cannot exclude that other cell types (e.g. fibroblasts, endothelial) contribute to the EMT-like signature we observed in the OE heart, however we present compelling evidence that this is mostly attributed to the CM population.

We demonstrated the involvement of YAP signaling downstream to ERBB2 in CMs, and its requirement for various ERBB2-related phenotypes. We obtained accumulating evidence that YAP activity is regulated by mechanotransduction signaling driven by ERBB2 activation. We present evidence that YAP activation occurs in the context of, and in association with, the unique altered cytoskeletons that are remodeled downstream to ERBB2 signaling. The altered mechanosensory state, and the biology it triggers, is not the typical output of ERBB2 signal relay. Being a receptor tyrosine kinase signaling amplifier in many types of cancer, the prescribed roles of ERBB2 revolve mostly around phosphorylation and robust recruitment of downstream mitogenic cascades (Moasser, 2007)(Wang and Xu, 2019)(Tzahar et al., 1996). In this study, we uncovered a novel role of ERBB2 as a master cytoskeleton regulator, which is a key organelle for CMs. The precise signal relay leading to the cytoskeletal reprogramming is out of the scope of this article, but it is clear that the changes transcend the actin cytoskeleton (Reischauer et al., 2014) and profoundly implicate other cytoskeletal structures like microtubules and intermediate filaments.

YAP binding partners associated with cytoskeletal and nuclear envelope compartments, force sensors upregulation and nuclear shearing all suggest that ERBB2 signals to YAP is mediated via cytoskeleton to the nucleus. Although key studies in the field of YAP mechanical regulation focus on actomyosin-driven mechanical relay (Dupont et al., 2011)(Aragona et al., 2013)(Elosegui-Artola et al., 2017)(Nardone et al., 2017)(Qiao et al., 2017)(Shiu et al., 2018), our Co-IP assay detected novel associations of YAP with the cardiac embryonic intermediate filament Nestin and other sarcomeric modulators. We postulate that because CMs have unique cytoskeletal structures (e.g the sarcomere is a highly specialized acto-myosin organelle), the regulators triggering mechanical YAP activation may differ from non-muscle cells. The commonality of the different mechanisms, seems to converge in the nucleus, which is the main mechanosensing structure of in the cell (Kirby and Lammerding, 2018). YAP activation via mechanosensory pathways is abrogated in case of perturbed nuclear envelope structure (Shiu et al., 2018) or perturbed cytoskeleton-to-nucleus force transfer (Elosegui-Artola et al., 2017)(Driscoll et al., 2015)(Lombardi et al., 2011). Mutations in nuclear envelope proteins, particularly LaminA/C, are strongly implicated in cardiac myopathies thus highlighting the susceptibility of CMs to nuclear envelope malformations (Sullivan et al., 1999)(Schreiber and Kennedy, 2013).

In line with our findings, pYAP S274 and pYAP S352 were detected (together or alone) in several reports as being phosphorylated during mitosis (Yang et al., 2013)(Bui et al., 2016) (Zhao et al., 2014)(Olsen et al., 2010). YAP phosphorylation on S274 and S352 peaks at metaphase, aligning with reports suggesting CDK1 as a kinase for these sites upon mitosis (Yang et al., 2013)(Bui et al., 2016). We show that the phosphorylation at those sites is required for mitosis, and is conserved among other proliferative CM systems, as in the case of embryonic CMs or adult CMs derived from YAP 5SA mice (Monroe et al., 2019).

The diverse filamentous pattern of pYAP S274 staining we observed (in non-dividing CMs) suggests that this form of YAP activation is dynamic and possibly indicative of a unique mode of mechanical stimulation, which is a derivative of the abundant cytoskeletal changes that underlie ERBB2 signaling activation in CMs.

We summarize our findings in a proposed model (Figure 6). ERBB2 activation remodels the CM cytoskeleton towards migration and force transmission via upregulation of intermediate filaments (IF) and microtubule (MT) infrastructures, at the expanse of contractile functions. Equipped with new cytoskeletons, OE CMs can sense and interact with the ‘rejuvenated’ ECM. As a result of this altered mechanic state, YAP is activated in OE CMs via association with cytoskeletal compartments leading to phosphorylation on S274 and followed by nuclear accumulation and association with the nuclear envelope. In the nucleus YAP also gets phosphorylated on S352. YAP transcriptional activity in the nucleus promotes mitosis, and the phosphorylation on S274 and S352 peaks at metaphase and is required for mitosis. Owing to the altered mechanic state, and related to YAP activation, EMT-like processes involving migration and ECM remodeling are triggered in OE CMs, contributing to the potent regenerative outcome of ERBB2 activation.

**Figure 6:**
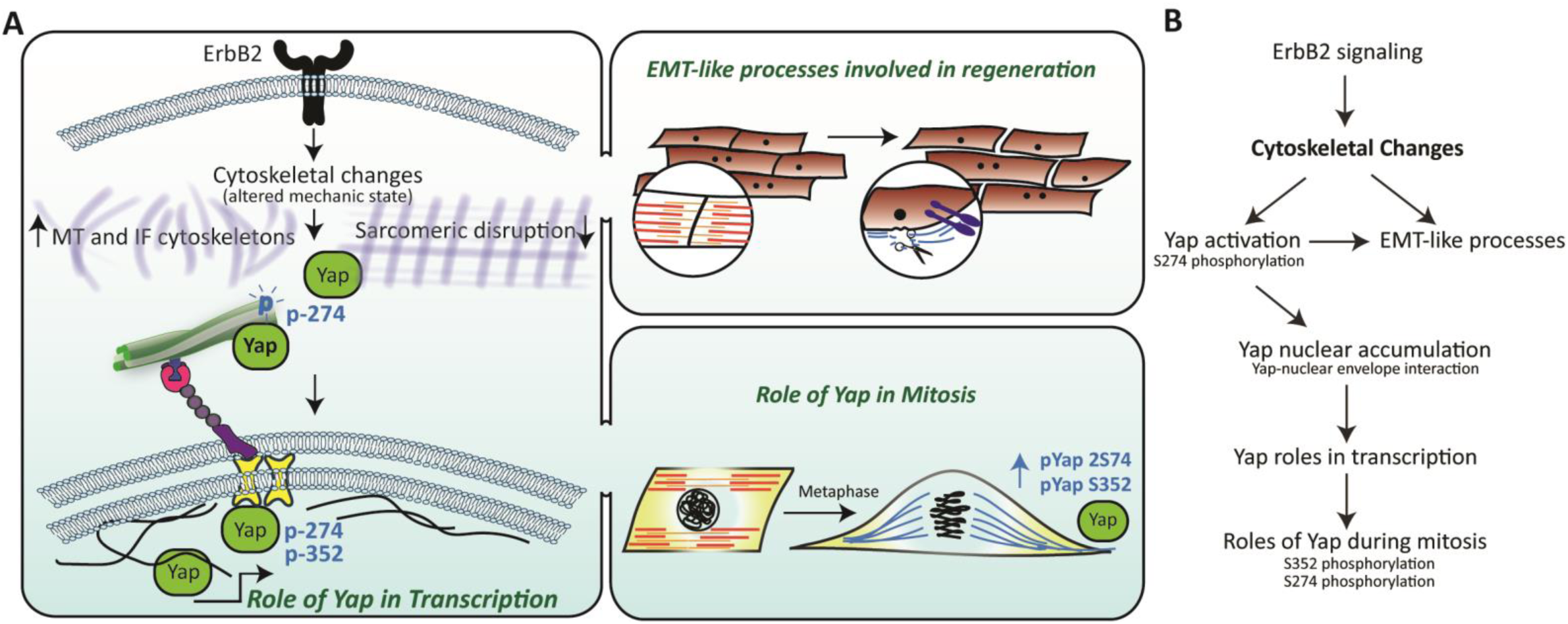
Proposed model for ERBB2-driven Yap induction and EMT-like processes. (A) Illustration: Upon ERBB2 activation in CMs, the cytoskeletal profile shifts from contractile specialization towards migration and force transmission via upregulation of intermediate filaments (IF) and microtubule (MT) infrastructures. Influenced by the altered mechanic state, Yap is activated in CMs via association with cytoskeletal compartments leading to phosphorylation on S274 and nuclear accumulation and association with the nuclear envelope, where it also gets phosphorylated on S352. Yap transcriptional activity in the nucleus promotes mitosis and EMT-like processes, and the phosphorylation on S274 and S352 peak at metaphase and are required for it. (B) Flow chart of (A).

## Author contributions

A.A. and E.T. conceived and designed the experiments. A.A with help from A. Shakked carried out most of the experiments and analyzed the data. Specifically, A. Shakked helped with animal studies, contributed to tissue culture work, RT-PCR, and spearheaded cloning of YAP mutants into AAV viruses and all revolving work and analysis. K.B.U. helped with animal studies, RNA - seq preparation and analysis and enabled the simultaneous RNA, protein and phosphoprotein acquisition method for HTS used throughout this study. A. Savidor and Y.L performed all proteomic analysis. D.K. and D.L. performed myocardial infarction experiments and echocardiographic analysis. O-Y.R helped with functional gelatin degradation assays and migration time-lapse microscopy. Y.M helped with 5SA heart sections. J.D. provided custom made and pYAP S274 antibody. B.G and J.F.M contributed to the planning and progression of the project. E.T supervised the entire project. A.A and E.T wrote the manuscript with editing contributions from all the authors.

## Acknowledgments

This study has been supported by grants to E.T from the European Research Council (ERC StG #281289, CM turnover, and ERC AdG #788194, CardHeal), the U.S.-Israel Binational Science Foundation (BSF) to both E.T. and J.F.M., the Israel Science Foundation (ISF), Foundation Leducq Transatlantic Network of Excellence, and Minerva foundation with funding from the Federal German Ministry for Education and Research. This work was supported by grants from the National Institutes of Health (HL127717, HL130804, HL118761 (J.F.M.); Vivian L. Smith Foundation (J.F.M.), State of Texas funding (J.F.M.). We thank Dr. Oded Singer for help with AAV preps and shared materials, Dr. Gilgi Friedlander for RNAseq analysis and inputs, and Noam Priel for teaching ‘trackmate’ software usage.

## Materials and Methods

### Mice

Animal experiments were approved by the Weizmann Institute of Science IACUC committee (Approval 13240419-3). CM-restricted, Doxycycline-inducible overexpression of ca*Erbb2* was generated as before (D’Uva et al., 2015). Doxycycline (Dox, TD02503, Harlan Laboratories) was administered in the food to repress transgene expression. For P7 cultures of WT/OE hearts, mice were additionally crossed with the αMHC-cre (Agah et al., 1997) and ROSA26-tdTomato (Madisen et al., 2010) transgenes in order to endogenously tag the CM lineage with tdTomato fluorescent protein. For the OE-KD mouse, inducible-Cre (αMHC-MerCreMer, 005657, Jackson) (Sohal et al., 2001) mice were crossed with LoxP *Yap* mice (027929, Jackson) (Zhang et al., 2010) and then αMHC-tTA (Yu et al., 1996) in one mouse, while TetRE-caErbB2 ca*Erbb2* (Xie et al., 1999) was crossed with LoxP *Yap* mice and ROSA26::tdTomato in another mouse, culminating with their cross with one another to generate OE-KD mice upon inheritance of all desired transgenes, and OE/WT littermates upon partial inheritance of transgenes. Genotyping primers are in Supplementary table 1.

### Myocardial infarction

Mice were sedated with isoflurane (Abbot laboratories) and were artificially ventilated following tracheal intubation. Experimental myocardial infarction was induced by permanent ligation of the left anterior descending coronary artery (LAD ligation) as described before (D’Uva et al., 2015). Following the closure of the thoracic wall, mice were injected s.c with Buprenorphin (0.066mg/Kg) as analgestic and were warmed for several minutes until recovery.

### Echocardiography parameters

Cardiac function was evaluated by transthoracic echocardiography performed on sedated mice with isoflourane (Abbot laboratories) using Vevo3100 (VisualSonics, Canada) in the approximate pulse range between 300-400bpm, and were analysed in the Vevo Lab 3.2.0 software (VisualSonics, Canada). Measurements for ejection fraction, systolic volume, diastolic volume, cardiac output, and stroke volume were calculated from the long axis. Measurements for LVAW and LVPW were calculated from the short axis, representing the thickness of the wall in the papillary plane.

For the delayed induction of ca*Erbb2* after injury (HF model, Figure 1), mice were approximately 3 months old in the beginning of the experiment and were randomized for sex. MI Injuries were performed as described above after individual baseline measurements of cardiac function. Injury severity was validated individually for every mouse, and only mice that lost at least 10%, but no more than 60% (at 2-3 weeks echocardiographic evaluation) of their baseline ejection fraction value entered the study. This is because injuries lesser then 10% were considered insufficient, and for injuries surpassing 60% function loss, ERBB2 signaling was insufficient in inducing regeneration.

### Sample preparation for HT assays

Adult mice approximately 3 months old were randomized for sex and subjected to MI (or Sham) as described above. Mice were injured and had Dox removal at the same day (Accute ErbB2 activation). 2 weeks post injury and Dox withdrawal, whole hearts were collected from 3 independent experimental cohorts, containing at least 4 biological repeats for every group. Hearts were briefly perfused with ice cold PBS and flash frozen in liquid Nitrogen. Using a mortar and pestle, frozen hearts were crushed to fine powder (intermittently maintaining nitrogen cooling) and were split to 3-ways for different processing methods for RNAseq, Proteomics and phosphoproteomics.

### RNA - seq

Working on ice, RNA was extracted and purified from the powder (see sample preparation section above) using miRNeasy kit (217004, Qiagen), according to the manufacturer instructions. Truseq libraries were made and barcoded, and subsequently subjected to 60bp single read solexa illumine sequencing, using the Solexa HiSeq. For RNA analysis, Adapters were trimmed using the cutadapt tool (Martin, 2011). Following adapter removal, reads that were shorter than 40 nucleotides were discarded (cutadapt option –m 40). Reads were sub-sampled randomly to have 23M Reads. TopHat (v2.0.10) was used to align the reads to the mouse genome (mm10)(Kim et al., 2013). Counting reads on mm10 RefSeq genes (downloaded from igenomes) was done with HTSeq-count (version 0.6.1p1) (Anders et al., 2013). Differential expression analysis was performed using DESeq2 (1.6.3)(Anders et al., 2013)(Love et al., 2014). Raw P values were adjusted for multiple testing using the procedure of Benjamini and Hochberg.

### Bioinformatic analysis

Ingenuity Pathway Analysis tool (https://apps.ingenuity.com/) was used for GO annotation of RNA seq and proteome data, using differential expression thresholds (as stated in the figure legend). The gene set enrichment analysis (GSEA) software (Subramanian et al., 2005) was used to perform gene set enrichment analysis on groups of genes, using the Hallmark (H) module. Heat maps were constructed for log transformed values using Gene-E software. For the values of each row, the row mean was subtracted and then divided by the standard deviation, and the color coding was done in a relative way for that row. Go term clustering was done in PANTHER software by using an overrepresentation test (GO Ontology database Release 2018-02-02), with a significance threshold of 0.05 for adjusted q-value.

### Proteomics, phosphoproteomics (processing, LC, MS, analysis)

Working on ice, protein was extracted from the powder (see ‘sample preparation’) in 600µl of SDT buffer (4%(w/v) SDS, 100mM Tris/HCl pH 7.6, 0.1M DTT) supplemented with phosphatase (Sigma, P0044) and protease (Dyn diagnostics, 11836170001) inhibitors via thorough homogenization. Samples were subjected to in-solution tryptic digestion using a modified Filter Aided Sample Preparation protocol (FASP)(Wisniewski et al., 2009). Tissue was further lysed by bead beating in the presence of SDT buffer supplemented with the inhibitors. Lysate was centrifuged at 16,000 g for 10min. 3 mg total protein were mixed with 2 mL Urea Buffer (UB) (8 M urea (Sigma, U5128) in 0.1 M Tris/HCl pH 8.0 and 50mM ammonium bicarbonate), loaded onto 30 kDa molecular weight cutoff filters (Sartorius VS15RH22) and centrifuged. 1.5 ml of UB were added to the filter unit and centrifuged at 14,000g for 40 min. Proteins were alkylated using iodoacetamide (10 mM final concentration) and washed twice with ammonium bicarbonate. Trypsin was then added (50:1 protein amount:trypsin) and samples incubated at 37°C overnight, followed by a second trypsin digestion for 4 hours at 37°C. Digested peptides were then collected into a clean tube by centrifugation, acidified with trifloroacetic acid, desalted using HBL Oasis (Waters 094225), speed-vac to dryness and stored in −80°C until analysis.

Liquid chromatography: ULC/MS grade solvents were used for all chromatographic steps. For protein expression, each sample was fractionated offline using high pH reversed phase followed by online low pH reversed phase separation. 200µg digested protein was loaded using high Performance Liquid Chromatography (Agilent 1260 uHPLC). Mobile phase was: A) 20mM ammonium formate pH 10.0, B) acetonitrile. Peptides were separated on an XBridge C18 column (3×100mm, Waters) using the following gradient: 3% B for 2 minutes, linear gradient to 40% B in 50min, 5 min to 95% B, maintained at 95% B for 5 min and then back to initial conditions. Peptides were fractionated into 15 fractions. The fractions were then pooled: 1 with 8, 2 with 9, 3 with 10, 4 with 11, 5 with 12, 6 with 13 and 7 with 14-15. Each fraction was dried in a speedvac, then reconstituted in 25 µL in 97:3 acetonitrile:water+0.1% formic acid. Each pooled fraction was then loaded and analyzed using split-less nano-Ultra Performance Liquid Chromatography (10 kpsi nanoAcquity; Waters, Milford, MA, USA). The mobile phase was: A) H2O + 0.1% formic acid and B) acetonitrile + 0.1% formic acid. Desalting of the samples was performed online using a Symmetry C18 reversed-phase trapping column (180 µm internal diameter, 20 mm length, 5 µm particle size; Waters). The peptides were then separated using a T3 HSS nano-column (75 µm internal diameter, 250 mm length, 1.8 µm particle size; Waters) at 0.35 µL/min. Peptides were eluted from the column into the mass spectrometer using the following gradient: 4% to 20%B in 105 min, 20% to 90%B in 5 min, maintained at 90% for 5 min and then back to initial conditions.

For phosphoproteomics, phosphopeptides were enriched from 1mg total protein digest using a ProPac IMAC-10 column (4×50mm) (Thermo Scientific, P.N. 063276). Loading buffer (A)-0.1% TFA, 30% ACN. Elution buffer (B)-0.3% NH4OH, pH 11.7. Peptides were loaded on the column in 10min with flow 0.1ml/min (0% B). Phosphopeptides were eluted from the column using the following gradient: first elution from 10 to 15 min with flow 0.6ml/min (B from 0% to 15%), second elution from 15min to 44 min with flow 0.02 ml/min(B from 15% to 30%), washing from 44 min to 50 min with flow 1ml/min (B from 30% to 50%) and following equilibration from 50min to 60min with flow 1 ml/min (0%B). The phosphopeptides containing peak (second elution fraction) was collected in 1.5 ml reaction vessels (∼1,2ml), speed-vac to dryness, and reconstituted in 25 µL in 97:3 acetonitrile:water+0.1% formic acid.

For LC-MS/MS analysis, phosphopeptides were separated online by low pH reversed phase liquid chromatography as described above with the following gradient: 4% to 20%B in 140 min, 20% to 90%B in 25 min, maintained at 90% for 5 min and then back to initial conditions.

Mass Spectrometry: The nanoUPLC was coupled online through a nanoESI emitter (10 μm tip; New Objective; Woburn, MA, USA) to a quadrupole orbitrap mass spectrometer (Q Exactive Plus, Thermo Scientific) using a FlexIon nanospray apparatus (Proxeon). For protein expression, data was acquired in DDA mode, using a Top10 method. MS1 resolution was set to 70,000 (at 400m/z) and maximum injection time was set to 20msec. MS2 resolution was set to 17,500 and maximum injection time of 60msec. For phosphoproeomics, data was acquired in DDA mode, using a Top20 method. MS1 resolution was set to 70,000 (at 400m/z) and maximum injection time was set to 100msec. MS2 resolution was set to 17,500 and maximum injection time of 120msec.

Data analysis: For protein expression, raw data was imported into the Expressionist® software (Genedata) and processed as described here (Shalit et al., 2015). The software was used for retention time alignment and peak detection of precursor peptides. A master peak list was generated from all MS/MS events and sent for database searching using Mascot v2.5 (Matrix Sciences). Data was searched against the mouse protein database downloaded from UniprotKB (http://www.uniprot.org/) appended with rat ErbB2 (P06494) sequence (caErbB2 transgene used in this study is from Rat) and 125 common laboratory contaminant proteins. Fixed modification was set to carbamidomethylation of cysteines and variable modifications were set to oxidation of methionines and deamidation of N or Q. Search results were then filtered using the PeptideProphet (Nesvizhskii et al., 2003) algorithm to achieve maximum false discovery rate of 1% at the protein level. Peptide identifications were imported back to Expressionist to annotate identified peaks. Quantification of proteins from the peptide data was performed using an in-house script (Shalit et al., 2015).

Data was normalized based on the total ion current. Protein abundance was obtained by summing the three most intense, unique peptides per protein. A Student’s t-Test, after logarithmic transformation, was used to identify significant differences across the biological replica. Fold changes were calculated based on the ratio of arithmetic means of the experimental groups.

For phosphoproteomics raw data was analyzed using MaxQuant_1.5.2. (Cox, Jurgen, Mann, 2008). Enzyme specificity was set to trypsin and up to two missed cleavages were allowed. Fixed modification was set to carbamidomethylation of cysteines and variable modifications were set to oxidation of methionines, deamidation of N or Q, and phosphorylation of S, T, or Y. Peptide precursor ions were searched with a maximum mass deviation of 4.5 ppm and fragment ions with a maximum mass deviation of 20 ppm. Peptide, protein and site identifications were filtered at an FDR of 1% using the decoy database strategy (MaxQuant’s “Reward” module). The minimal peptide length was 7 amino-acids and the minimum Andromeda score for modified peptides was 40. Peptide identifications were propagated across samples using the match-between-runs option checked. Searches were performed with the label-free quantification option selected. Fold changes were calculated based on the ratio of arithmetic means of the experimental groups. A Student’s t-Test, after logarithmic transformation, was used to identify significant differences across the biological replica.

### Co-Immunoprecipitation (processing, LC, MS, analysis)

Co-IP was done using magnetic beads Co-IP kit (Universal Magnetic CO-IP Kit, Active Motif, 54001-am) according to the manufacturer’s instructions. Briefly, adult hearts were excised, quickly perfused and washed in ice-cold PBS cold and homogenized (Omni TH-02) in freshly made lysis buffer. 1000µg of the soluble fraction were taken for IP using 2µg of YAP antibody (NB110-58358, Novus) or IgG control (#I5006, Sigma), and 25µl of magnetic beads, in a total of 500µl Co-IP buffer made fresh the same day and incubated on the rotor at 4C over night. Beads were pulled aside with the magnet and washed with freshly made wash buffer, as instructed. If samples were subsequently sent of Mass Spectromery, they were washed in the wash buffer only twice, followed by additional 2 washes in PBS supplemented with protease inhibitors (1:100, Sigma), and straight away were processed for Mass Spec in the following manner. Samples were subjected to on-bead tryptic digestion. Briefly, washed beads after co-IP were suspended in 8M urea, 0.1M Tris HCl buffer, pH 7.6, and incubated at room temp for 30 min. Proteins were then reduced with 100 mM DTT (5mM final conc) at room temp for 1 hr and then alkylated using iodoacetamide (10 mM final concentration) for 45 min in the dark at room temp. Urea concentration was diluted 5-fold by addition of 50mM ammonium bicarbonate. Trypsin was then added (50:1 protein amount:trypsin) and samples incubated at 37°C overnight, followed by a second trypsin digestion (100:1 protein amount:trypsin) for 4 hours at 37°C. Digested peptides were then collected by centrifugation of the beads and transferring the supernatant into a clean tube. Peptides were acidified with trifloroacetic acid, desalted using HBL Oasis (Waters 094225), speed vac to dryness and stored in −80°C until analysis.

Liquid chromatography: digested proteins were loaded and analyzed using split-less nano-Ultra Performance Liquid Chromatography (10 kpsi nanoAcquity; Waters, Milford, MA, USA). The mobile phase was: A) H2O + 0.1% formic acid and B) acetonitrile + 0.1% formic acid. Desalting of the samples was performed online using a Symmetry C18 reversed-phase trapping column (180 µm internal diameter, 20 mm length, 5 µm particle size; Waters). The peptides were then separated using a T3 HSS nano-column (75 µm internal diameter, 250 mm length, 1.8 µm particle size; Waters) at 0.35 µL/min. Peptides were eluted from the column into the mass spectrometer using the following gradient: 4% to 25%B in 50 min, 25% to 90%B in 5 min, maintained at 90% for 5 min and then back to initial conditions.

Mass Spectrometry: the nanoUPLC was coupled online through a nanoESI emitter (10 μm tip; New Objective; Woburn, MA, USA) to a quadrupole orbitrap mass spectrometer (Q Exactive HF, Thermo Scientific) using a FlexIon nanospray apparatus (Proxeon). Data was acquired in data dependent acquisition (DDA) mode, using a Top10 method. MS1 resolution was set to 120,000 (at 400m/z) and maximum injection time was set to 60msec. MS2 resolution was set to 15,000 and maximum injection time of 60msec.

Data analysis: raw data was analyzed using MaxQuant_1.6.0.16.). The raw data was searched against the mouse protein database downloaded from UniprotKB (http://www.uniprot.org/) appended with rat ErbB2 (P06494) sequence (caErbB2 transgene used in this study is from Rat) and 125 common laboratory contaminant proteins. Enzyme specificity was set to trypsin and up to two missed cleavages were allowed. Fixed modification was set to carbamidomethylation of cysteines and variable modifications were set to oxidation of methionines, protein N-terminal acetylation, and phosphorylation of S, T, or Y. Peptide precursor ions were searched with a maximum mass deviation of 4.5 ppm and fragment ions with a maximum mass deviation of 20 ppm. Peptide, protein and site identifications were filtered at an FDR of 1% using the decoy database strategy. The minimal peptide length was 7 amino-acids and the minimum Andromeda score for modified peptides was 40. Peptide identifications were propagated across samples using the match-between-runs option checked. Searches were performed with the label-free quantification option selected. Fold changes were calculated based on the ratio of arithmetic means of the experimental groups. A Student’s t-Test, after logarithmic transformation, was used to identify significant differences between experimental groups across the biological replica. Dataset was filtered for differentially detected (Pvalue ≤0.05, fold change≥±2) binding partners in the IP reactions for OE/WT, while discarding as background binding partners that were also differentially detected in the corresponding IgG fractions.

### qPCR

RNA from whole hearts was isolated using the NucleoSpin kit (740955.50, Machery Nagel) according to the manufacturer’s instructions. A High Capacity cDNA Reverse transcription kit (Applied Biosystems, 4374966) was used to reverse transcribe 1µg of purified RNA according to the manufacturer’s instructions. qPCR reactions were performed using Fast SYBR Green PCR Master Mix (4385614, Thermo Fischer Scientific). Oligonucleotide sequences for real-time PCR analysis performed in this study are listed in Supplementary Table1.

### Western blot

Total cell lysates were isolated with RIPA buffer supplemented with 1:100 protease (P8340, Sigma) and 1:100 phosphatase inhibitor cocktails (P5726 Sigma and P0044 Sigma). Protein concentrations were quantified by BCA assay (catalog, Thermo Fisher). Lysates was separated in denaturating Tris Glycine gels and wet transferred (in the case of proteins of interest above 100kD) or semi-dry transferred (in the case of proteins of interest below 100kD) onto 0.44micron PVDF membranes. Membranes were blocked and probed for antibodies described in Supplementary table 2.

### Cell Culture

Primary cardiac cultures were isolated from P7 mice using a neonatal dissociation kit (Miltenyi Biotec,130-098-373) using the gentleMACS homogenizer, according to the manufacturer’s instructions and cultured in Gelatin-coated (0.1%, G1393, Sigma) wells with DMEM/F12 (D6421, Sigma) medium supplemented with L-glutamine (1%, 03-020-1B, Biological Industries), Na-pyruvate (1%, 03-042-1B, Biological Industries), nonessential amino acids (1%, 01-340-1B, Biological Industries), penicillin, streptomycin, amphothericin (1%, 03-033-1B, Biological Industries), horse serum (5%, 04-004-1A, Biological Industries) and FBS (10%, 04-007-1, Biological Industries) (‘complete-medium’) at 37◦C and 5% CO2 for 24h. Afterwards, medium was replaced with FBS-depleted medium (otherwise same composition) for additional 48h after which the cultures were fixated and stained (for basic IF analysis). For gelatin degradation, migration and virus infection assays, P7 cardiac culture isolation was started the same and adjusted as specified below.

### Functional Geltain degradation assay

Fluorescently labelled gelatin (0.2 mg porcine skin gelatin; #G2500, Sigma) was prepared by Alexa Fluor™ 488 labeling Kit (A10235, molecular probes, Thermo Fisher Scientific), according to the manufacturer’s instructions. Glass-bottomed 96-well plates were coated, with ratios of 1:10 labeled gelatin (0.2%): non-labeled gelatin (0.2%). For degradation assays, P7 cardiac cultures were seeded (see cell culture section above) at low density (aprx 500-700 CMs/well) in ‘complete medium’ and after 48h the wells were fixated and stained for cTNT (CM marker), phalloidin (P1951, Sigma) and DAPI (see immunofluorescence section). Wells were randomly imaged using DeltaVision microscope using a 40x/0.75 air objective and degradation area was assessed by ImageJ software. As OE CMs can move, this assay was relatively short lived (fixation 48 hours after seeding) and the degradation area was attributed to any particular cell only if it was found specifically beneath that cell (and not in the vicinity), hence the sparse seeding.

### Migration assay

Ibidy inserts (80209, Ibidi) were tightly positioned, each in a separate well in a 24 well plate, that has been coated with 1:40 (in PBS) fibronectin coating (03-090-1, Biological Industries) and allowed to dry in order to allow tight adhesion of the insert. P7 WT/OE cardiac cultures were seeded in the 2 sided on an ibidy chamber in ‘complete medium’ and allowed to settle for 24 hours. After 24 hours, chamber has been removed using sterile forceps and the medium changed to fresh ‘complete medium’ for additional 4 days to allow migration, following fixation and staining.

### Time lapse movie and single-cell tracking

P7 WT/OE cardiac cultures tagged with tdTomato fluorescent protein were seeded on fibronectin (03-090-1, Biological Industries) coated 1:40 (in PBS) wells in ‘complete medium’. After 24 hours, the medium has been changed to fresh ‘complete medium’ and time lapse imaging has been set up. Movies were taken for period of 96 hours, using DeltaVision microscope using a 10x air objective, at imaging intervals of 10 minutes. Single-cell migration quantification was done using ImageJ software with the Trackmate Add-on. Individial CMs were tracked based on cells being present for at least 75% of the time in the field.

### Immunofluorescence

For P7 cardiac culture: Cardiac cultures were fixated with 4% PFA for 10 minutes in room temperature on the shaker, following permeabilization with 0.5% Triton X-100 in PBS for 5min, and blocking with 5% bovine serum albumin (BSA) in PBS containing 0.1% Triton for 1h at room temperature. Primary antibody solution was prepared in the blocking solution (detailed list in Supplementary table 2), and incubated either for 1h at RT or over night at 4c, while shaking. The cultures were subsequently washed 3 times in PBS for 10 minutes while shaking, and incubated for 1h with the appropriate secondary antibodies (abcam or Jackson) followed by 10 min incubation with DAPI for DNA visualization. Finally, the cultures were washed with PBS additional 3 times and were imaged on the DeltaVision, Ti2 Nikon or Olympus fluorescent light microscopes.

For heart sections: Hearts were briefly perfused in-situ with 4% PFA, extracted and washed in cold PBS, following additional fixation at 4c o/n.while shaking in 4% PFA-filled tube. Then, hearts were by paraffin embedded and sectioned. Slides were de-paraffinized and antigen retrival was performed using EDTA/Citrate buffer, following gradual coolling of 1h while shaking at RT. Then, sections were permeabilized for 5 minutes with 0.5% triton, and blocked for at least 1h following application of primary ab in blocking solution for o/n incubation at 4°C. Importantly, section perimeters were thoroughly traced with a hydrophobic pap-pen to allow complete and even coverage of the section under a thick pool of primary ab. Slides were then thoroughly washed for 3 times, while shaking in RT, following the application of secondary Abs (Jackson, Abcam) for 1h at RT. Finally, DAPI was applied to counterstain nuclei. Slides were additionally washed and mounted with Immumount (9990402, Thermo Scientific) and imaged on the Ti2 Nikon, Olympus or Ziess Confocal spinning disc microscopes. In the cases of staining for YAP, pYAP S274, pYAP S352 (only in sections) and Sun2, HRP-based tyramide amplification has been used, according to manufacturer’s instructions (B40925, Thermo scientific). Antibodies used for IF can be found in Supplementary table 2.

### pYAP S274 and S352 antibodies

pYAP S274 rabbit polyclonal custom-made antibody was provided by the Dong lab (Yang et al., 2013). pYAP S352 rabbit polyclonal custom-made antibody was generated by Sigma using the peptide GTPNAVS**S**PGMSQEL (phosphorylated serine in bold) in a double affinity purification procedure.

### YAP phosphomutant cloning and AAV infection

The mouse YAP1 WT gene with a Myc-tag in the pCMV6 backbone was purchased from Origene (MR226049). YAP S274A and YAP S352A mutants were generated by PCR assembly. All primers described below for mutant cloning can be found in Supplementary table 1. For YAP S274A, a 910-bp fragment was amplified with YAP1-Str-F1-F & YAP1-274A-F1-R and a 708-bp fragment was amplified with YAP1-274A-F2-F and YAP1-End-F2-R from the pCMV6-YAP1 WT template. Assembly PCR was performed with YAP1-Str-F1-F and YAP1-End-F2-R using the 910- and 708-bp fragments as a template followed by ligation into pCMV6. The same process was used to create the YAP S352A, using the 352A-unique primers. To create pCMV6-YAP2A, pCMV6-YAPS274A and pCMV6-YAPS352A were restricted to generate a vector containing the S274A mutation and an insert containing the S352A mutation, which were subsequently ligated. All pCMV6-YAP1 variants were amplified with YAP1-Str-F1-F and YAP1-End-SalI and ligated into pAAV-CMV plasmid (kindly provided by Dr. Oded Singer, Weizmann Institute Viral Core). The YAP5SA gene was amplified with primers YAP1-5SA-Str-BglII and YAP1-5SA-End-SalI using pQCXIH-Myc-YAP-5SA (Addgene #33093) as a template and ligated into the pAAV-CMV plasmid. All pAAV-CMV-YAP variants were validated by DNA sequencing and were used to generate crude AAV-DJ preparations. P7 WT/OE cardiac cultures were isolated as described above and seeded in ‘complete medium’. 24 hours after seeding, each well was treated with 2.5E8vg of virus diluted in 200ul of FBS-depleted media for 48 hours, after which the cells were fixated and stained for markers of proliferation.

### Scar quantification

Scar quantification was based on Masson’s Trichrome staining done on representative serial cardiac sections spanning all the heart. In each section, the % of scar (blue) has been denoted by the angle of fibrotic tissue (in degrees, out of 360) as measured with a protractor (using the middle of the heart as the center operating under the assumption circularity of the cardiac section). The % of scarring was then averaged between all sections of a heart spanning from the beginning of the ventricles downwards to the apex.

### Statistics

Sample size was chosen empirically following previous experience in the assessment of experimental variability. Generally, all experiments were carried out with n≥3 biological replicates (except for the Mass Spec analysis on cut-out bands that was done only once (Figure S7F)). The analyzed animal numbers or cells per groups are described in the respective figure legends. We ensured that experimental groups were balanced in terms of animal age, sex and weight unless otherwise specified. Animals were genotyped before the experiment and were caged together and treated in the same way. Statistical analysis was carried out using Prism software. Whenever comparing between 2 conditions, data was analyzed with two tailed student’s t-test (except for in Figure 1G, where one tailed t-test had been used). If comparing more than 2 conditions, we employed ANOVA analysis with multiple comparisons. In bar plots comprised of colour coded dots (such as in Figure1S) the different shades differentiate data points derived from different biological repeats to demonstrate the distribution of data across different experiments. The statistical analysis is derived from the average value of all data points from a particular biological repeat of an experiment, eventually comparing biological repeats (and not their individual constituents). In all figures, measurements are reported as mean, and the error bars denote s.e.m. Throughout the study, threshold for statistical significance was considered for p-values≤0.05, denoted by one asterisk (∗) if P ≤0.05, two (∗∗) if P <0.01, three (∗∗∗) if P <0.001 and four (∗∗∗∗) if P <0.0001 as stated in the figure legend.

