## Supplementary material for "ERBB2 drives YAP activation and EMT-like processes during cardiac regeneration": supp data

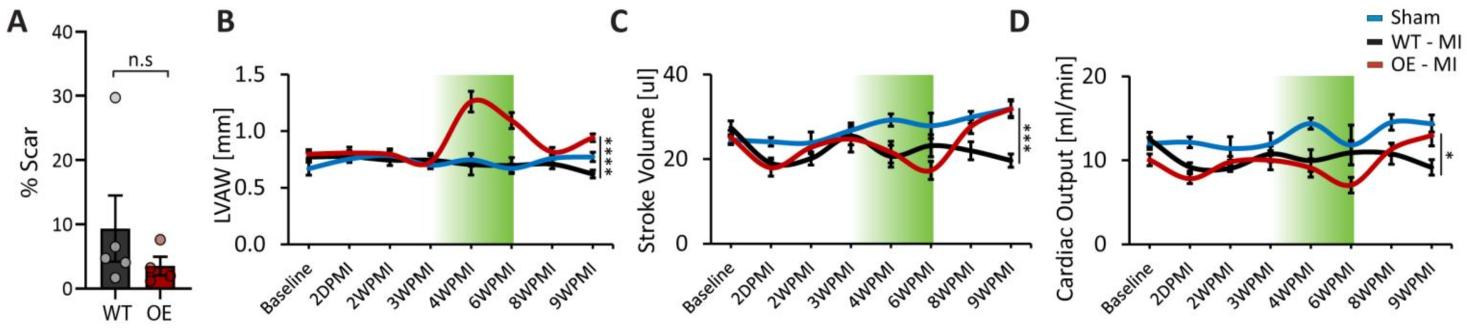


**Figure S1: Delayed and transient ERBB2 induction promotes functional and anatomic improvement in the heart, Related to Figure 1.**

(A) % Scar quantification for the time point of 3WPMI (Figure 1A, orange arrow) n=5 for WT, n=4 for OE. (B- D) Cardiac parameters derived from echocardiographic analysis. n=11 mice for OE, n=16 mice for WT, n=5 for Sham; (B) LVAW thickness. (C) Stroke volume (D) Cardiac output.

* $p < 0.05$; *** $p < 0.001$; **** $p<0.0001$. Error bars indicate SEM. All experiments were performed for at least 3 biological repeats.

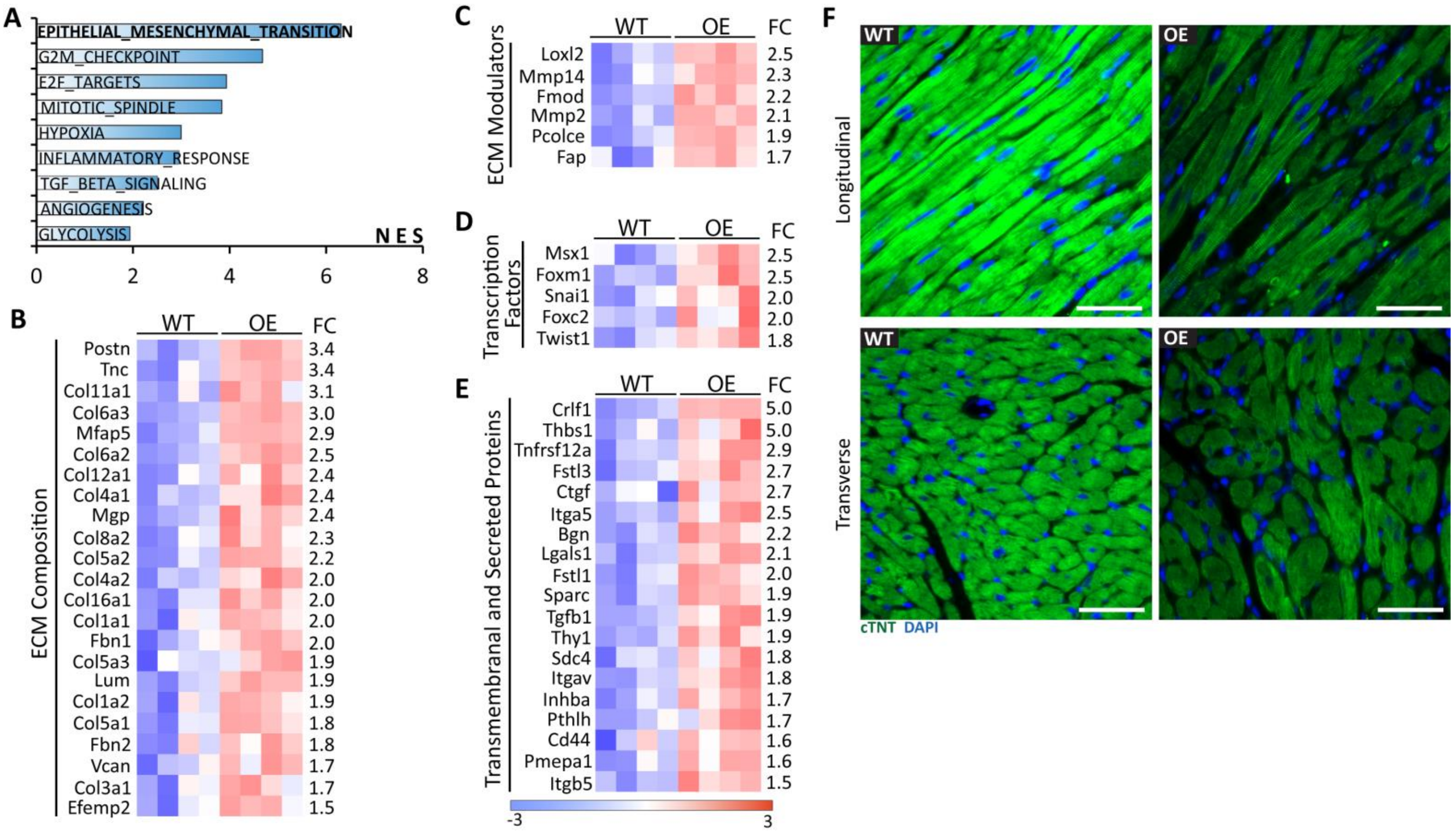


**Figure S2: EMT hallmarks are upregulated in OE hearts, Related to Figure 1.**

(A) Bulk RNA- seq WT-MI and OE-MI analysed using GSEA hallmark module (relates to Figure 1H). Bar plots are depicting the normalized enrichment scores (NES). (B-E) RNA- seq derived heat map of the indicated genes belonging to the EMT hallmark (as shown in (A) and Figure 1H). Genes were annotated as (B) ECM composition (C) ECM modulators (D) Transcription factors and (E) Transmembranal and secreted proteins. FC to the right indicates fold change of the detected transcripts by RNA- seq. All displayed results are statistically significant at adjusted *pv* ≤0.05. n=4 for WT, n=4 for OE. (F) IF of indicated proteins in WT/OE hearts**.** Scale bar, 50 µm.

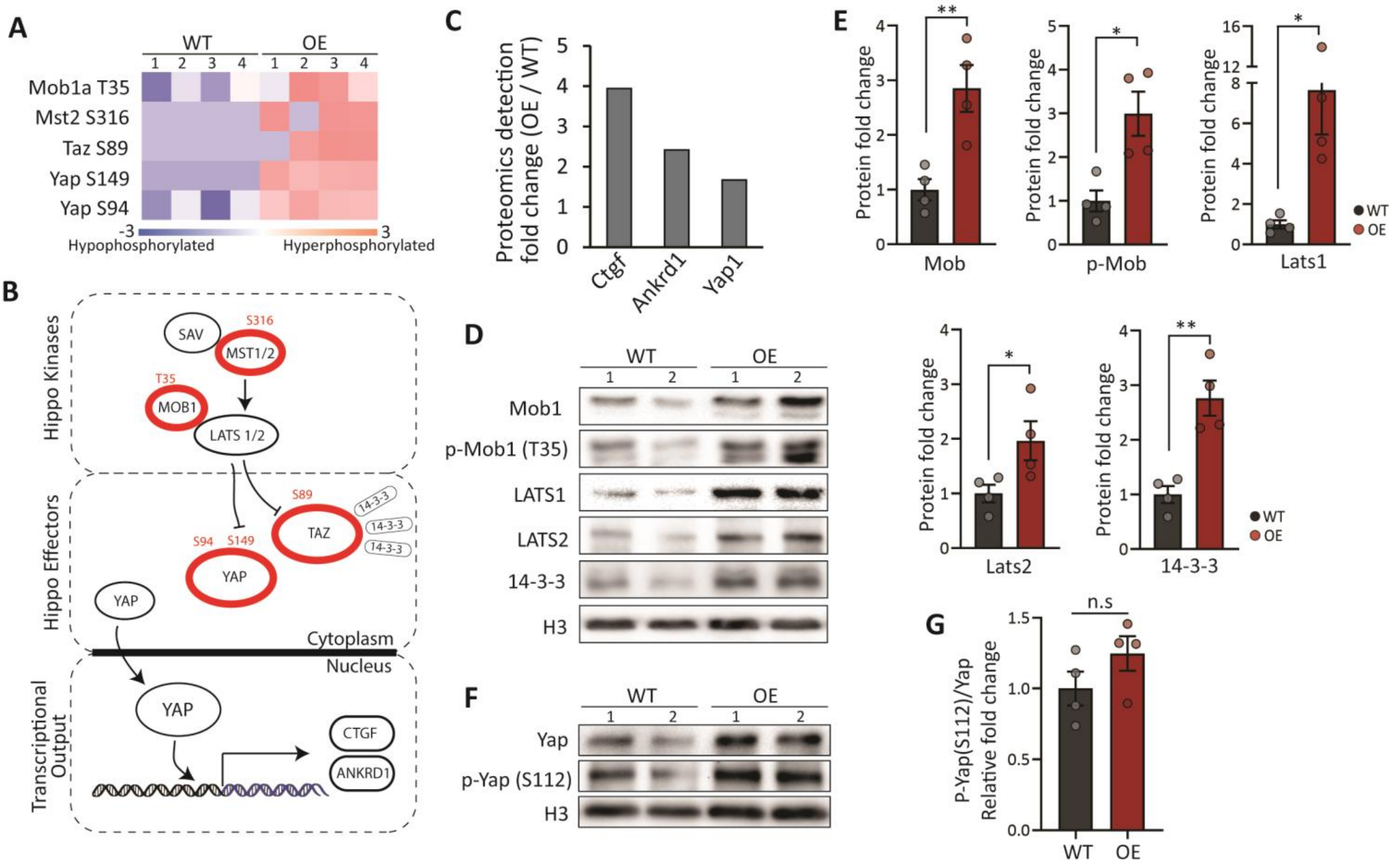


**Figure S3: Hippo pathway is not attenuated in Erbb2 OE hearts, Related to Figure 2.**

(A) Heat map of Phosphoproteomic analysis of the indicated proteins belonging to the Hippo pathway (See also Figure 2A). Phosphorylation site indicated to the left. n=4 for WT, n=4 for OE. All displayed results are at *pv* < 0.05. (B) A scheme of the Hippo pathway representing the phosphoproteomic data in (A) by a red circle with the corresponding site. (C) Proteomic data of OE/WT fold change for Yap and the targets CTGF and ANKRD1 (n=4 for WT, n=4 for OE). (D) WB analysis of the indicated proteins from in vivo adult WT/OE heart lysates. (E) Quantification of (D) (n=4 for WT and n=4 for OE). (F) WB analysis of the indicated proteins from in vivo adult WT/OE heart lysates. (G) Quantification of (F) (n=4 for WT and n=4 for OE).

* $p < 0.05$; ** $p < 0.01$; Error bars indicate SEM. All experiments were performed for at least 3 biological repeats.

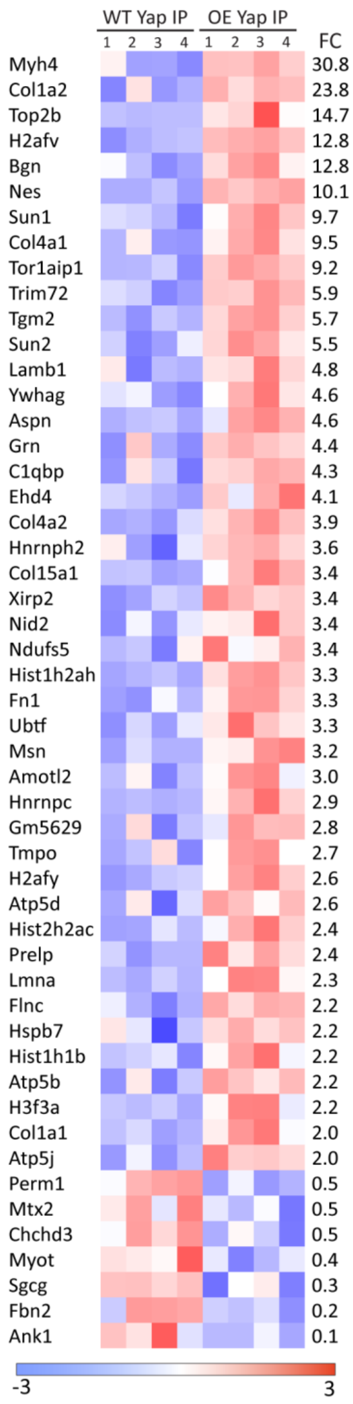


**Figure S4: Yap CO-IP-MS full binding partner list, Related to Figure 4**

Heat map of Yap binding partners of the assay in Figure 4A. FC to the right indicates fold change between OE to WT IP reactions of the detected protein by Mass Spec. All displayed results are at pv $\leq 0.05$.

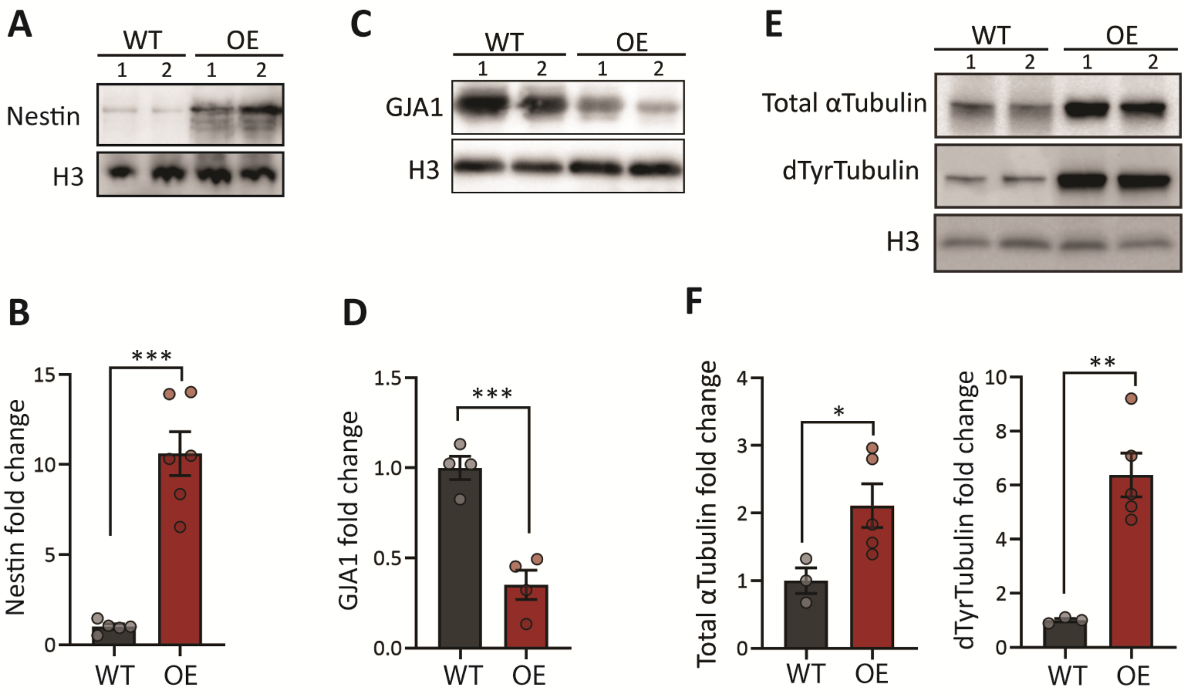


**Figure S5: Cytoskeletal changes in OE hearts, Related to Figure 4**

(A) WB analysis of the indicated proteins from adult WT/OE heart lysates. (B) Quantification of (A) (n=5 for WT, n=6 for OE). (C) WB analysis of the indicated proteins from adult WT/OE heart lysates. (D) Quantification of (C) (n=4 for WT, n=4 for OE). (E) WB analysis of the indicated proteins from adult WT/OE heart lysates. (F) Quantification of (E) (n=3 for WT and n=5 for OE).

* $p$ < 0.05; ** $p$ < 0.01; *** $p$ < 0.00. Error bars indicate SEM.

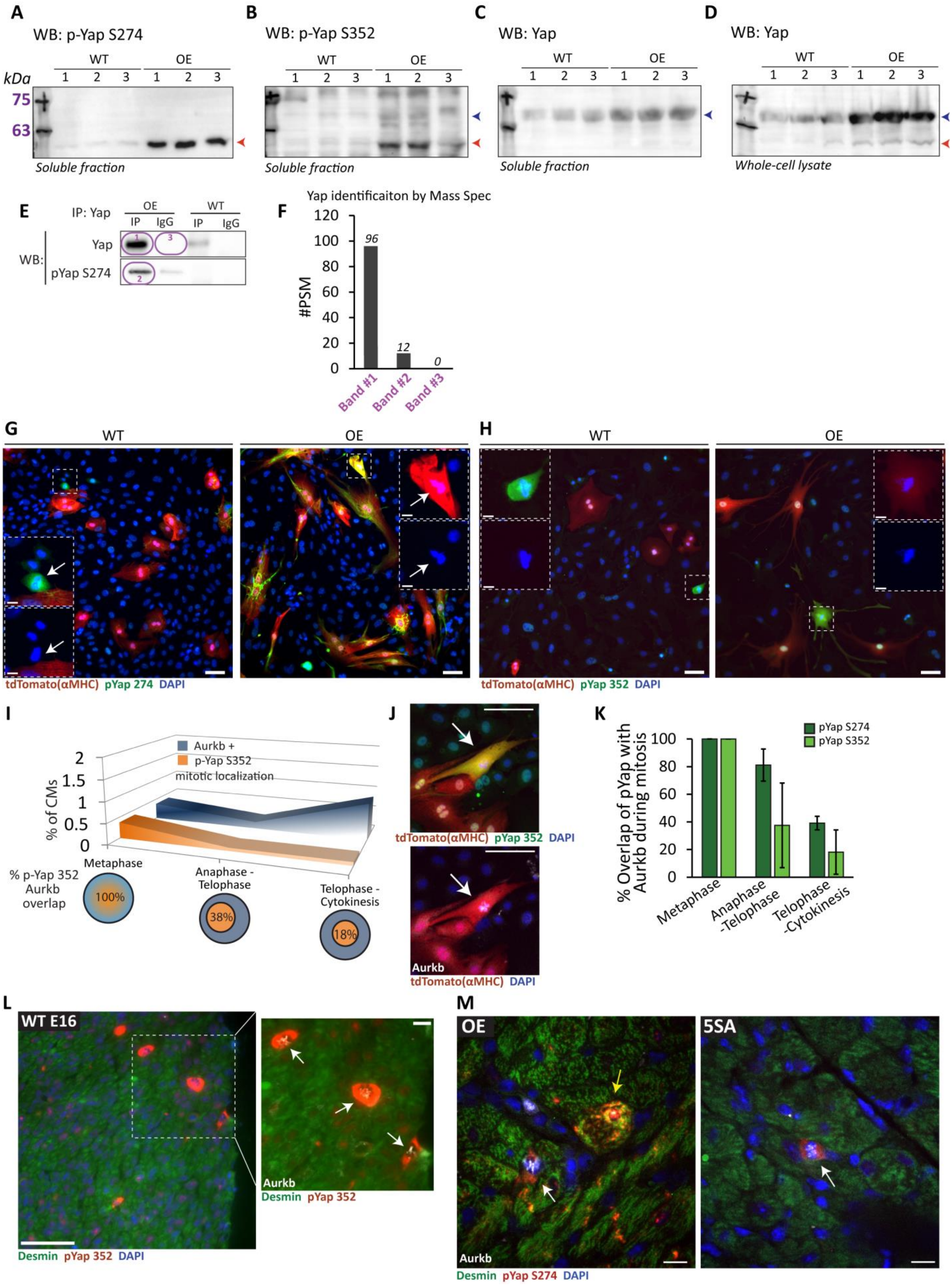
A
WB: p-Yap S274
kDa
75
63
WT
OE
1 2 3 1 2 3
Soluble fraction
B
WB: p-Yap S352
Soluble fraction
C
WB: Yap
Soluble fraction
D
WB: Yap
Whole-cell lysate
E
IP: Yap
OE
WT
IP IgG IP IgG
WB:
Yap
pYap S274
F
Yap identificaiton by Mass Spec
#PSM
96
12
0
Band #1
Band #2
Band #3
G
WT
OE
tdTomato(αMHC) pYap 274 DAPI
H
WT
OE
tdTomato(αMHC) pYap 352 DAPI
I
Aurkb +
p-Yap S352
mitotic localization
% of CMs
Metaphase
Anaphase - Telophase
Telophase - Cytokinesis
% p-Yap 352 Aurkb overlap
100%
38%
18%
J
tdTomato(αMHC) pYap 352 DAPI
Aurkb
tdTomato(αMHC) DAPI
K
pYap S274
pYap S352
% Overlap of pYap with Aurkb during mitosis
Metaphase
Anaphase -Telophase
Telophase -Cytokinesis
L
WT E16
Aurkb
Desmin pYap 352
Desmin pYap 352 DAPI
M
OE
5SA
Aurkb
Desmin pYap S274 DAPI

**Figure S6: Phosphorylated Yap on S274 and S352 is prevalent in OE hearts, Related to Figure 5**

(A) WB analysis of p-Yap S274 on the soluble fraction of adult WT/OE heart lysates. Red arrow head points to the band below 63kDa (i.e the “lower” band). (B) WB analysis of p-Yap S352 on the soluble fraction of adult WT/OE heart lysates. Blue arrow head points to the band between 75kDa and 63kDa (i.e the “upper” band), and the red arrow head points to the “lower” band. (C) WB analysis of Yap on the soluble fraction of adult WT/OE heart lysates. Blue arrow head points to the “upper” band. (D) WB analysis of Yap on whole cell lysates (include cytoskeletal components that are usually underrepresented in soluble extracts) of adult WT/OE hearts. Blue and red arrow heads point to both “upper” and “lower” bands. (E) Yap IP (and IgG control) from OE/WT heart lysates blotted for Yap and pYapS274. The purple circles represent cut-out bands that were analysed by Mass-Spec. (F) Bands represented in the purple circles in (E) were cut and analysed by mass Spectrometry. Bars represent the number of peptide-spectrum match (#PSM) detection of Yap protein for the corresponding band numbers. (G) Representative images of IF analysis for the indicated proteins of P7 WT/OE cardiac cultures. Scale bar 50µm. Inset in WT highlights with an arrow a non-CM cell at metaphase with peaking pYap S274 stain. Inset in OE highlights with an arrow a CM at metaphase with peaking pYap S274 stain. Scale bar 10 µm. (H) Representative images of immunofluorescence analysis for the indicated proteins of P7 WT/OE cardiac cultures. Scale bar 50µm. Inset in WT highlights with an arrow a non-CM cell at metaphase with peaking pYap S352 stain. Inset in OE highlights with an arrow a CM at metaphase with peaking pYap S352 stain. Scale bar 10 µm. For panels (G-H) CM lineage endogenously tagged with tdTomato. (I) Quantification of CMs positive for Aurkb (blue curve) and pYap S352 mitotic peak localization (orange curve) as counted for the different stages of mitosis plotted on the x axis. (n= 2274 CMs). Bottom Venn diagram show the degree of overlap between of pYap S352 mitotic peak localization from all Aurkb events. (J) Representative images of immunofluorescence analysis for the indicated proteins at metaphase P7 OE cardiac cultures (CM lineage endogenously tagged with tdTomato). Scale bar 50µm. (K) Bar plot depicting the degree of overlap of pYap S274 (dark green) and pYap S352 with the different stages of mitosis plotted on the x axis. This is summary of data from panel (I) and from Figure 5D. (L) IF of embryonic WT E16.5 heart sections for the indicated proteins. Scale bar 50µm. The inset to the right shows the selected area counterstained with Aurkb. Arrows points to CMs in metaphase. Scale bar 10µm. (M) IF of adult heart sections for the indicated proteins for OE and 5SA. White arrows point to a CM in metaphase and the yellow arrow points to a pre-metaphase (aurora is still nuclear) CM. Scale bar, 10μm. All experiments were performed for at least 3 biological repeats, except for band analysis which was done once.

**Supplementary table 1:** Primers

**qRT – PCR Primers**

| Gene | Forward (5' to 3' sequence) | Reverse (5' to 3' sequence) |
|---|---|---|
| *Anln* | GTTAAAACTCGAATGCAAAGGCT | AGTTGGCACTGGTGCAAAGTA |
| *Ccnb1* | TGCGAACCAGAGGTGGAACTGGA | TTCCATTGGGCTTGGAGAGGGAGT |
| *Cdk6* | GGCGTACCCACAGAAACCATA | AGGTAAGGGCCATCTGAAAACT |
| *Cyr61* | CTGCGCTAAACAACTCAACGA | GCAGATCCCTTTCAGAGCGG |
| *Enah* | CTGGTGGCTCAACTGGGTTC | TGCCCACAACTCTGAATGTGT |
| *Sntb1* | AACAGGCAGCTAGAAATTCACTC | AAGTCACCAGCGTTGGAATGA |
| *Ankrd1* | TGCGATGAGTATAAACGGACG | GTGGATTCAAGCATATCTCGGAA |
| *Aurkb* | CAGAAGGAGAACGCCTACCC | GAGAGCAAGCGCAGATGTC |
| *Birc5* | GAGGCTGGCTTCATCCACTG | CTTTTTGCTTGTTGTTGGTCTCC |
| *Ctgf* | GAGGAAAACATTAAGAAGGGCAAA | CGGCACAGGTCTTGATGA |
| *Sphk1* | GGAGGAGGCAGAGATAACCTT | GACCCAACTCCTCTGCACACA |
| *Aspm* | TGGCTATGAGTGAATGCTCTTCC | TCGCGTAAAAACAGTGGCAAG |
| *Bub1b* | GAGGCGAGTGAAGCCATGT | TCCAGAGTAAAAGCGGATTTCAG |
| *Cdc2* | TTTCGGCCTTGCCAGAGCGTT | GTGGAGTAGCGAGCCGAGCC |
| *Foxm1* | CAGAATGCCCCGAGTGAAACA | GTGGGGTGGTTGATAATCTTGAT |
| *Plk1* | CTTCGCCAAATGCTTCGAGAT | TAGGCTGCGGTGAATTGAGAT |
| *Rhamm* | CCTTGCTTGCTTCGGCTAAAA | CTGCTGCATTGAGCTTTGCT |
| *Mmp2* | TTCTGGTCAAGGTCACCTGTC | CAAGTTCCCCGGCGATGTC |
| *Mmp14* | AGCACTGGGTGTTTGACGAA | GTCTTCCCATTGGGCATCCA |

**Genotyping Primers**

| Gene | Forward (5' to 3' sequence) | Reverse (5' to 3' sequence) |
|---|---|---|
| *TetRE-caErbb2* | AAGAAGAGCCCAAGCTGGA | GTGTACGGTGGGAGGCCTAT |
| *αMHC-tTA* | CGCTGTGGGGCATTTTACTTTAG | CATGTCCAGATCGAAATCGTC |
| *αMHC-Cre* | GGCCAGCTAAACATGCTTCA | ACACCAGAGACGGAAATCCATC |
| *Mer Cre Mer* | TCTATTGCACACAGCAATCCA | CCAGCATTGTGAGAACAA GG |
| *Yap flox* | AGGACAGCCAGGACTACACAG | CACCAGCCTTTAAATTGAGAAC |
| *Rosa26-tdTomato* | GGCATTAAAGCAGCGTATCC | CTGTTCCTGTACGGCATGG |

**Primers for the generation of phospho-Yap mutants**

| **Primer name** | **5' to 3' sequence** |
|---|---|
| Yap1-Str-F1-F | GTAATACGACTCACTATAGGGCG |
| Yap1-End-F2-R | GCTGCCAGATCCTCTTCTGAG |
| Yap1-274A-F1-R | CTCCCTGTGGGGCCTGGGGAGCCAAGGGTGG |
| Yap1-274A-F2-F | GGCTCCCCAGGCCCCACAGGGAGGCGTCC |
| Yap1-352A-F1-R | GAGACATCCCAGGAGCAGACACTGCATTCGGAGTCCC |
| Yap1-352A-F2-F | GAATGCAGTGTCTGCTCCTGGGATGTCTCAGGAATTG |
| Yap1-End-*Sal*I | CGAGCGTCGACTTAAACCTTATCGTCGTCATCC |
| Yap1-5SA-Str-*Bgl*II | TTAGAGATCTGAACGGTGCATTGGAACGGACC |
| Yap1-5SA-End-*Sal*I | CGAGCGTCGACCTATAACCATGTAAGAAAGCTT |

**Supplementary Table 2:** Antibodies

**IF antibodies**

| Antibody against | Manufacturer | CAT # | Concentration |
|---|---|---|---|
| cTNT | Abcam | ab33589 | 1:200 |
| cTNI | Abcam | ab47003 | 1:200 |
| Desmin | Santa Cruz | SC-23879 | 1:100 |
| Vimentin | Abcam | ab24525 | 1:200 |
| Sun2 | Abcam | ab124916 | 1:200 |
| Ki67 | Cell Marque | 275R-14 | 1:200 |
| Aurora | BD Transduction Laboratories | BD 611082 | 1:200 |
| PH3 | Cell Signaling | 9701 | 1:200 |
| Yap | Novus Biologicals | NB110-58358 | 1:150 |
| 274 | (Yang et al., 2013) | custom ab | 1:400 |
| 352 | Sigma | custom ab | 1:200 |
| Nestin | Abcam | ab11306 | 1:200 |
| Tub | Cell Signaling | 3873 | 1:100 |
| dTyrTub | Abcam | ab48389 | 1:100 |
| GJA1 | Abcam | ab11370 | 1:200 |

**WB antibodies**

| Antibody against | Manufacturer | CAT # | Concentration |
|---|---|---|---|
| Yap | Novus Biologicals | NB110-58358 | 1:2000 |
| pYap S112 | Cell Signaling | 13008 | 1:1000 |
| Mob1 | Cell Signaling | 3863 | 1:1000 |
| p-Mob1 | Cell Signaling | 8699 | 1:1000 |
| LATS1 | Cell Signaling | 3477 | 1:500 |
| LATS2 | Novus Biologicals | NB200-199 | 1:5000 |
| 14-3-3 | Cell Signaling | 8312 | 1:1000 |
| ErbB2 | Cell Signaling | 2165 | 1:1000 |
| H3 | Abcam | ab1791 | 1:5000 |
| pLAMNA | Cell Signaling | 13448S | 1:1000 |
| LMNA | Santa Cruz | SC-7293 | 1:200 |
| Sun2 | Abcam | ab124916 | 1:5000 |
| Tub | Cell Signaling | 3873 | 1:1000 |
| dTyrTub | Abcam | ab48389 | 1:1000 |
| GJA1 | Abcam | ab11370 | 1:8000 |
| Nestin | Abcam | ab11306 | 1:1000 |
| 274 | (Yang et al., 2013) | custom ab | 1:2000 |
| 352 | Sigma | custom ab | 1:2000 |

Yang, S., Zhang, L., Liu, M., Chong, R., Ding, S.J., Chen, Y., and Dong, J. (2013). CDK1 phosphorylation of YAP promotes mitotic defects and cell motility and is essential for neoplastic transformation. Cancer Res. *73*, 6722–6733.
